# Persister state-directed transitioning and vulnerability in melanoma

**DOI:** 10.1101/2020.04.01.999847

**Authors:** Heike Chauvistré, Batool Shannan, Sheena M. Daignault, Robert J. Ju, Daniel Picard, Stefanie Egetemaier, Renáta Váraljai, Antonio Sechi, Farnusch Kaschani, Oliver Keminer, Samantha J. Stehbens, Qin Liu, Xiangfan Yin, Kirujan Jeyakumar, Felix C. E. Vogel, Clemens Krepler, Vito W. Rebecca, Linda Kubat, Smiths S Lueong, Jan Forster, Susanne Horn, Marc Remke, Michael Ehrmann, Annette Paschen, Jürgen C. Becker, Iris Helfrich, Daniel Rauh, Markus Kaiser, Sheraz Gul, Meenhard Herlyn, José Neptuno Rodríguez-López, Nikolas K. Haass, Dirk Schadendorf, Alexander Roesch

## Abstract

Melanoma is a highly plastic tumor characterized by dynamic interconversion of different cell identities depending on the biological context. For example, melanoma cells with high expression of the H3K4 demethylase KDM5B (JARID1B) rest in a slow-cycling, yet reversible persister state. Over time, KDM5B^high^ cells can promote rapid tumor repopulation with equilibrated KDM5B expression heterogeneity. The cellular identity of KDM5B^high^ persister cells has not been studied so far, missing an important cell state-directed treatment opportunity in melanoma. Here, we have established a doxycycline-titratable system for genetic induction of permanent intratumor expression of KDM5B and screened for chemical agents that phenocopy this effect. Transcriptional profiling and cell functional assays confirmed that the dihydropyridine phenoxyethyl 4-(2-fluorophenyl)-2,7,7-trimethyl-5-oxo-1,4,5,6,7,8-hexa-hydro-quinoline-3-carboxylate (termed Cpd1) supports high KDM5B expression and directs melanoma cells towards differentiation along the melanocytic lineage and to cell cycle-arrest. The high KDM5B state additionally prevents cell proliferation through negative regulation of cytokinetic abscission. Moreover, treatment with Cpd1 promoted the expression of the melanocyte-specific tyrosinase gene specifically sensitizing melanoma cells for the tyrosinase-processed antifolate prodrug 3-O-(3,4,5-trimethoxybenzoyl)-(-)-epicatechin (TMECG). In summary, our study provides proof-of-concept for a new dual hit strategy in melanoma, in which persister state-directed transitioning limits tumor growth and plasticity and primes melanoma cells towards lineage-specific elimination.

## Introduction

Cellular plasticity describes the capacity of cells to switch from one phenotype to another and is considered a major driver of non-genetic tumor evolution. Recent time-resolved transcriptomic profiling points to an understanding particularly of melanoma as a heterogeneous ecosystem that dynamically transitions through variably differentiated persister states to escape from exogenous pressure such as debulking therapies ^1–3^.

One hallmark of persister cells across different cancers is that they transiently rest in a chromatin-mediated slow-cycling state characterized by a high, but reversible nuclear expression of histone H3 lysine demethylases (KDMs) ^4–6^. Other features of persister cells are maintenance of embryonic survival or metabolic programs balancing their dependency on oxidative mitochondrial respiration and detoxification of oxygen and lipid radicals ^5, 7, 8^. To date, the majority of studies have focused on the selective elimination of persister cells as small subsets within the total tumor population by exploiting single vulnerabilities like fatty acid oxidation, autophagy, or ferroptosis ^9–11^. Previous approaches to selectively eliminate H3K4 demethylase expressing melanoma cells (KDM5B^high^), e.g. based on their biological dependence on oxidative ATP production, showed some effect, but suffered from the high metabolic flexibility of this tumor entity ^12, 13^. Surprisingly, the cellular identity of KDM5B^high^ melanoma persisters has not been unraveled so far, missing an important treatment opportunity. Moreover, strategies to manipulate the dynamics in phenotype switching of persister cells and to combine those with cell identity-specific elimination are lacking in the field.

Melanomas are usually characterized by a continuous expression spectrum of KDM5B^high^, KDM5B^intermediate^, and KDM5B^low^ cells. In comparison, benign melanocytic nevi express KDM5B at unexpectedly high levels across the majority of cells ^14^. Single-sorted melanoma cells can rapidly re-establish KDM5B heterogeneity irrespective of their initial KDM5B expression level ^15^. Even the pronounced enrichment for KDM5B^high^ states, which is typically seen under cytotoxic stress *in vitro* and *in vivo* ^5, 16, 17^, rapidly reverts to normal distribution in surviving melanoma cells. This suggests that slow-cycling KDM5B^high^ persister cells represent a transient source for tumor repopulation, but longevity of melanoma requires a dynamic KDM5B tumor composition.

All together, this indicates that the slow-cycling KDM5B^high^ persister state could have different, maybe even opposing effects on tumor fate depending on the biological context and might be exploited as a therapeutic target for new tumor elimination strategies in melanoma. In this study, we have systematically investigated (i) the actual differentiation phenotype of the KDM5B^high^ melanoma persister state, (ii) the consequences of forcing melanoma cells to ectopically express KDM5B at high levels (like nevi) without the chance to spontaneously revert to normal expression heterogeneity and (iii) whether KDM5B-directed cell state transitioning be used as a therapeutic vulnerability to eradicate melanoma cells.

## Results

### Models for enforced and persistent intratumor expression of KDM5B^high^ states

KDM5B expression in melanoma is usually heterogeneously distributed, whereas benign melanocytic nevi express KDM5B at high levels across the majority of cells (Fig. 1A ^15^). High KDM5B expression is significantly associated with increased melanoma fitness and poor patient survival (http://cancergenome.nih.gov/, Fig. 1B). Other cancer entities or other KDM5 family members show weaker or no association with survival (Fig. S1). To create an experimental model that allows the exogenous induction of KDM5B protein expression, we cloned a Tet-On 3G-system, in which KDM5B expression is driven by a doxycycline-inducible PTRE3G promoter, and established stable melanoma cell clones (WM3734*^Tet3G-KDM5B^* and WM3734*^Tet3G-EGFP^* control, Fig. S2A-C). After clonal expansion and doxycycline induction, we observed a dose-dependent upregulation of KDM5B mRNA and protein level (Fig. 1C, Fig. S2C), concomitant with an inverse decrease in H3K4me3 protein levels confirming the expression of functional KDM5B (Fig. S2D).

**Figure 1.**
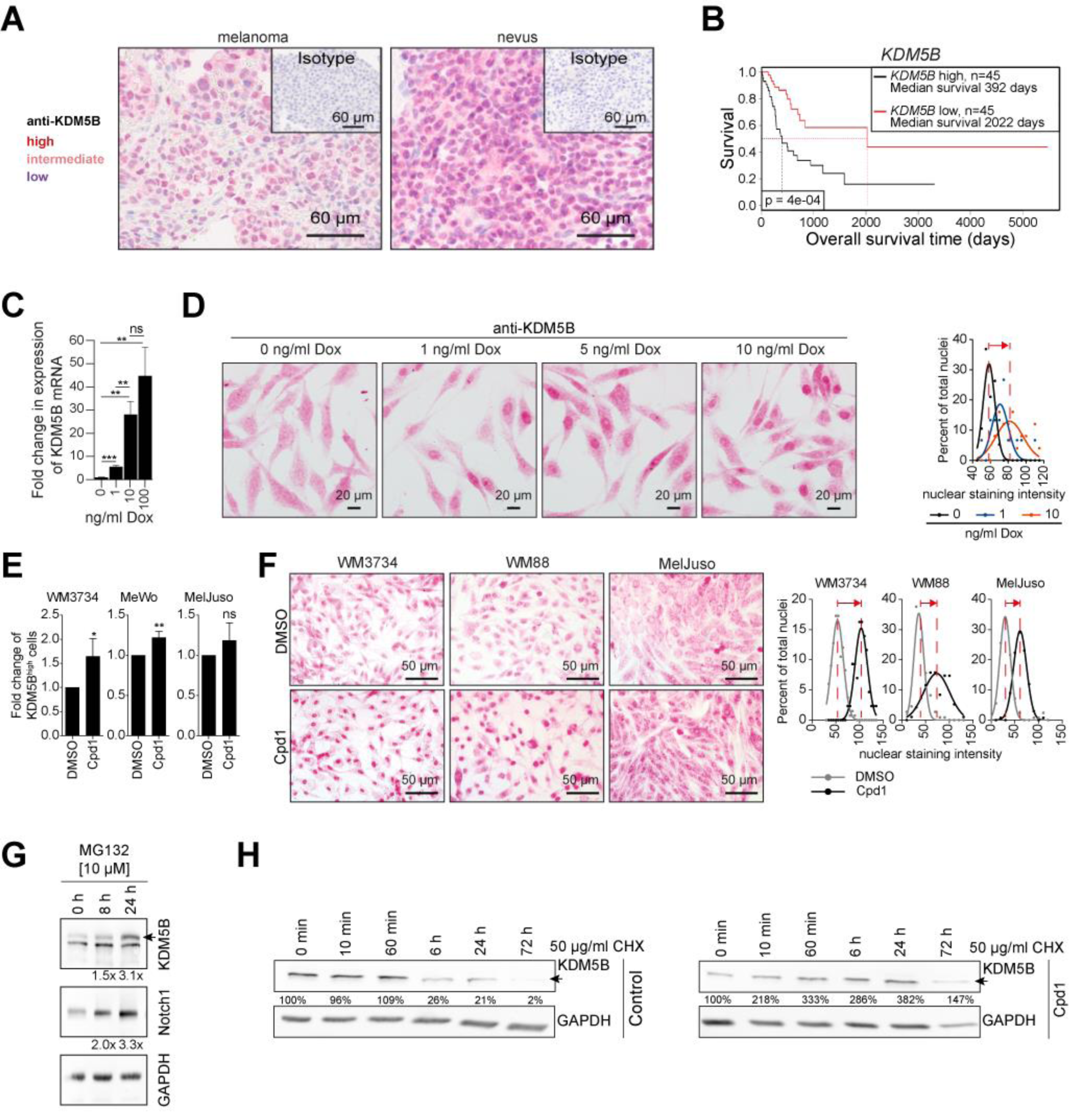
Expression modulation of the histone H3K4 demethylase KDM5B/JARID1B. (**A**) Anti-KDM5B immunostaining of a melanoma patient sample (left) compared to a benign human nevus (right). Highly positive nuclei are dark red, medium positive nuclei are light red, low expressing nuclei are blue. Isotype controls are shown in the upper right corners. Representative images of at least 5 different tissue samples. (**B**) Kaplan-Meier survival curves of cutaneous melanoma patients was calculated from the TCGA data set based on a 10% KDM5B expression threshold (10% highest expression, black, vs. 10% lowest, red, TCGA browser tool v0.9). (**C**) Quantitation of KDM5B mRNA induction after 24 h of doxycycline (Dox) treatment as assessed by qPCR. Mean ±SD; **P≤0.01, ***P≤0.001 by t-test. Shown is one representative example of two different clones. (**D**) Anti-KDM5B nuclear immunostaining of WM3734*^Tet3G-KDM5B^* cells after Dox-titration at the indicated concentrations for 24 h (left, representative images; right, quantitation shown as normalised frequency distribution of nuclear staining intensity). Shown is one representative out of 3 clones. **(E)** Flow cytometric detection of endogenous KDM5B protein levels after treatment with Cpd1 for 72 h. Shown are summarized data of 4 independent experiments. Mean ±SD; significance was tested by t-test; *P≤0.05, **P≤0.01. (**F**) Anti-KDM5B nuclear immunostaining of 3 different melanoma cell lines (WM3734, WM88, MelJuso) after 72 h of 10 µM Cpd1 treatment (left, representative pictures; right, quantitation shown as normalised frequency distribution of nuclear staining intensity). (**G** and **H**) Time course of KDM5B protein levels after treatment of WM3734 cells with MG132 (10 µM, **G**) or with Cpd1 (10 µM) plus cycloheximide (CHX, 50 µg/ml, n=2, **H**).

The KDM5B protein level was stably upregulated under continuous doxycycline treatment for at least 3 weeks (Fig. S2D). KDM5B immunostaining showed a shift towards increased nuclear signals across the majority of WM3734*^Tet3G-KDM5B^* cells (Fig. 1D). From here on, we denote this scenario as enforced KDM5B^high^ persister state.

To obtain a second, independent method for the modulation of KDM5B expression, which additionally allows simple application to several melanoma cell lines or *in vivo*, we developed a cell-based compound screening assay applying our previously published *KDM5B-promoter-EGFP-*reporter construct stably expressed in WM3734 melanoma cells [WM3734^KDM5Bprom-EGFP^ cells, ^15^, Fig. S3A]. This fluorescence-based model facilitated monitoring the dynamic nature of the KDM5B transcription state. After screening a 7,500 compound library, primary hits were counter-screened in a *CMV-promoter-EGFP*-reporter assay and confirmed hits were further validated in dose response curves and independent assays (Fig. S3B-D). The main criteria for hit compound selection were the absence of immediate overall cell toxicity at 72 h of treatment and modulation of the KDM5B-promoter-driven expression of EGFP (abbreviated K/EGFP, for more details see methods). Compounds that directly increased K/EGFP-signals were not considered because of possibly masking effects by passive enrichment for KDM5B^high^ cells as previously seen for various cytotoxic drugs ^5^. Instead, K/EGFP signal decrease was of particular interest because it could indirectly indicate a persistent increase of the endogenous KDM5B protein level as a result of chemical compound treatment (for example through a negative feedback of high KDM5B protein levels on mRNA transcription).

As confirmed by flow cytometric analysis and by immunocytology, the top hit compound 2-phenoxyethyl 4-(2-fluorophenyl)-2,7,7-trimethyl-5-oxo-1,4,5,6,7,8-hexahydroquinoline-3-carboxylate (PubChem name BAS00915510, here abbreviated Cpd1), which belongs to the dihydropyridine group of calcium channel inhibitors, reproducibly increased nuclear KDM5B protein levels irrespective of the genotype of the melanoma cell lines tested (Fig. 1E, F, Table S1). A structurally homologous compound that failed to change K/EGFP transcription and endogenous KDM5B protein levels was selected as negative control for subsequent experiments (abbreviated Neg4, Table S1, Fig. S3C, D). Quantitation of the KDM5B expression localization in nuclear vs. cytoplasmic compartments confirmed a Cpd1 time-dependent increase of specifically nuclear KDM5B (Fig. S3E) phenocopying the effect of Tet-On 3G-induced expression of KDM5B. Again, this was independent of the melanoma cell type tested (Fig. S3F).

Using the proteasome inhibitor MG132, we observed a steady increase of endogenous KDM5B protein after 8 and 24 h of treatment suggesting the proteasome as a likely degradation mechanism for endogenous KDM5B (Fig. 1G, Notch 1 used as positive control). To examine if KDM5B protein is better protected from degradation after Cpd1-mediated enrichment in the nucleus, we performed co-treatment with the protein synthesis inhibitor cycloheximide (Fig. 1H). Without Cpd1, KDM5B protein levels steadily declined with an almost complete loss after 72 h. The presence of Cpd1 prolonged this process suggesting that the enforced nuclear enrichment of KDM5B protects from protein degradation and, thus, may counteract the dynamic adjustment of KDM5B heterogeneity normally seen in melanoma.

### Enforced expression of KDM5B directs melanoma cells towards a slow-growing persister state

Standard MTT assays, caspase 3 and annexin V measurement, or 7AAD flow cytometry showed no major apoptotic or toxic effect in short-term Cpd1-treated melanoma cell lines and WM3734*^Tet3G-KDM5B^* cells (Fig. 2A, Fig. S4A, B). For Cpd1, a maximum feasible dose of 10 µM was assumed based on cross-cell line comparisons in MTT assays (Fig. S4C) and was used for all following experiments. In congruence with reports on KDM5B-associated therapy persistence ^5, 16, 17^, we observed decreased drug susceptibility upon KDM5B upregulation irrespective of the melanoma cell line tested (Fig. 2A).

**Figure 2.**
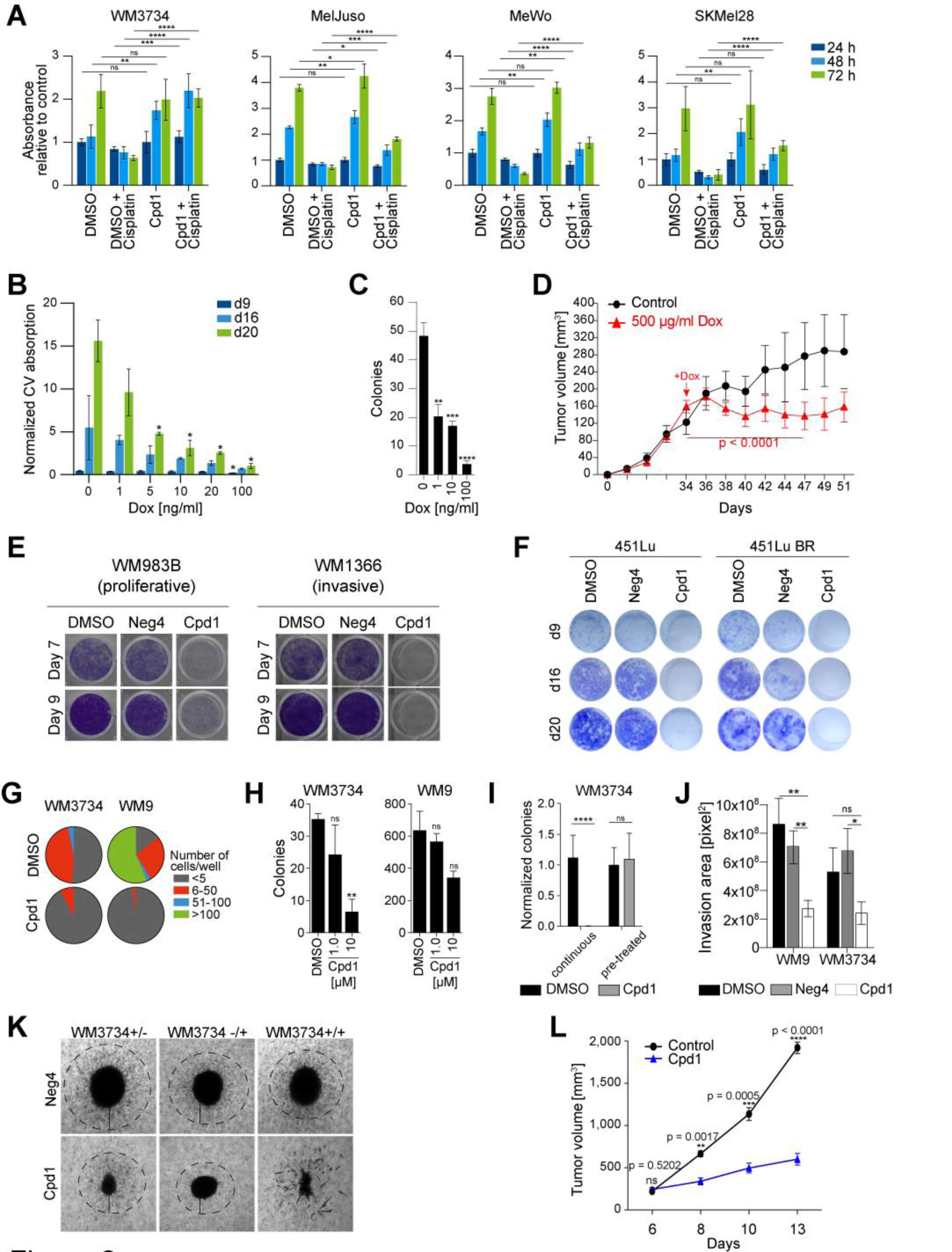
*In vitro* and *in vivo* effects of enforced KDM5B expression. (**A**) MTT cell viability assays of WM3734, MelJuso, MeWo and SKMel28 cells (representative example of n=3). Cpd1 treatment started at day -3, then 10 µM cisplatin was added as indicated. Readout was done after 24, 48, and 72 h. Data are shown as mean ±SD. Significance was determined by t-test (* P<0.05, **P≤0.01, ***P≤0.001, ****P≤0.0001). (**B**) Quantitation of long-term clonogenic growth after gradual KDM5B induction over 9, 16, and 20 days. Data are depicted as mean ±SD. Significance was calculated by t-test (* P<0.05). Shown is one out of 6 clones. (**C**) Soft agar colony formation after 52 days of KDM5B induction. Shown is one representative experiment out of three independent biological replicates. Data are depicted as mean ±SD; **P≤0.01, ***P≤0.001, ****P≤0.0001 (t-test). (**D**) Growth curves of WM3734*^Tet3G-KDM5B^* cells subcutaneously injected into nude mice. 500 µg/ml Dox was added to the drinking water starting from day 34. Data are shown as mean ±SEM with n=6 mice in the control and n=10 mice in treatment group. Significance was determined by a linear mixed-effect spline model. This data reflect one experiment. (**E**) Representative pictures of clonogenic growth assay of WM983B and WM1366 cells continuously treated over 7 and 9 days with 10 µM of Cpd1 or Neg4. Shown is one representative experiment out of 3 independent biological replicates. (**F**) Representative pictures of clonogenic growth assay of 451Lu and 451Lu BR cells treated over 9, 16 and 20 days with 10 µM of Cpd1 vs. DMSO or Neg4 controls. Shown is one out of 2 cell lines. (**G**) 2D colony formation capacity of single-seeded WM3734 and WM9 cells after continuous treatment with 10 µM Cpd1 for 19 days. Shown are means of three independent biological replicates. (**H**) Soft agar colony formation of WM3734 and WM9 cells under constant Cpd1 treatment at the indicated doses over 30 days and (**I**) compared to short-term treatment for 72 h before seeding without treatment continuation (for each cell line, experiments were performed once or twice in triplicate reaction, respectively). Mean ±SD; **P≤0.01, ****P≤0.0001 by t-test. (**J**) Quantitation of collagen invasion of WM3734 spheroids under 10 µM of Neg4 or Cpd1 at day 10. Shown is one representative experiment out of three independent biological replicates. Mean ±SD; *P≤0.05, **P≤0.01 by t-test. (**K**) Representative pictures of melanoma spheroids at day 10 after collagen-embedding. Neg4 vs. Cpd1 (10 µM) treatment was done either before collagen-embedding of cells (left, +/-) or after completed spheroid formation (middle, -/+) or before and after collagen-embedding (right, +/+). (**L**) Growth curves of murine CM melanoma cells subcutaneously injected into female C57BL/6N mice and treated with Cpd1. Data are shown as mean ±SEM with n=3/5 mice per treatment group. Significance was determined by t-test. This data reflect one experiment.

Next, we tested how melanoma cell populations behaved, when the KDM5B expression spectrum was constantly directed to a higher level without the chance to dynamically revert to normal heterogeneity. Enforced expression of KDM5B in the WM3734*^Tet3G-KDM5B^* model significantly reduced cell numbers in long-term 2D growth assays and decreased anchorage-independent 3D-colony formation in a dose-dependent manner (Fig. 2B, C, Fig. S4D). Growth of naïve WM3734 and WM3734*^Tet3G-EGFP^* control cells was not affected by doxycycline (Fig. S4E). The KDM5B-mediated effect on tumor proliferation was then studied *in vivo*. We allowed xenografted WM3734*^Tet3G-KDM5B^* cells to establish tumors up to approximately 150 mm^3^ before doxycycline was supplemented to the drinking water. This led to a significant and dose-dependent tumor growth delay over nearly two weeks (control group n=6, treatment group n=10, Fig. 2D; p<0.05, linear mixed-effect model).

Long-term treatment with Cpd1 significantly reduced the growth of different melanoma cell lines irrespective of their genotype and of whether those were allocated to the classic ‘invasive’ or ‘proliferative’ phenotype (Fig. 2E, Fig. S5A, B; Fig. S5C shows examples for treatment up to 16 days). Also melanoma cells that had previously developed resistance to MAPK inhibition responded to chemical enforcement of the KDM5B phenotype supporting the observation that chronically drug-resistant melanoma cell populations re-establish normal KDM5B heterogeneity after initial KDM5B enrichment (Fig. 2F, Fig. S5D-G).

In addition, limited dilution assays and 3D-colony formation indicated a loss of tumor repopulation properties of melanoma cells under constant exposure to Cpd1 (Fig. 2G, H). In contrast, pre-treatment of melanoma cells with Cpd1 for only 3 days before seeding onto soft agar was not sufficient to reduce the number of formed colonies again suggesting spontaneous reversibility of the KDM5B phenotype under normal conditions (Fig. 2I, Fig. S5H). Accordingly, growth arrest of melanoma cells usually seen under persistent exposure to Cpd1 was reversible after 1 week of compound removal (data not shown). Finally, continuous treatment with Cpd1 reduced invasive capacity of melanoma cells in a 3D-collagen spheroid model (Fig. 2J). Interestingly, Cpd1 strongly impaired the capacity of melanoma cells to form proper spheroids; especially when treatment commenced before collagen-embedding (Fig. 2K).

Finally, we tested Cpd1 in an immunocompetent syngeneic mouse melanoma model established previously ^18^. After confirmation of the phenotypic effects of Cpd1 treatment in murine melanoma cells *in vitro* (Fig. S5I, J), CM cells were subcutaneously injected into C57BL/6N mice and treated with Cpd1. This resulted in a significant anti-tumor growth effect in comparison to the control (Fig. 2L). *Ex vivo* immunostaining confirmed a Cpd1-induced shift towards high KDM5B expression states (Fig. S5K).

In sum, our results suggest that KDM5B expression dynamics can be limited by exogenous genetic or chemical manipulation providing a new possibility to therapeutically manipulate the slow-growing tumor phenotype.

### KDM5B-mediated cell cycle arrest and inhibition of cytokinetic abscission

Although persister cells have been repeatedly described to be slow-cycling in various cancer entities ^4, 6, 9^, their true cellular identity and proliferation phenotype remains elusive in melanoma. We now demonstrated by DNA content cell cycle analyses that enforced KDM5B expression dose-dependently increases the proportion of G0/1-arrested cells across various genetically different melanoma cell lines (n=5, Fig. 3A, Fig. S6A). Complimentary single cell cycle imaging of FUCCI-WM164 cells ^19^ treated with Cpd1 confirmed a significant delay in G1 phase progression in a majority of tracked cells and an increase in cell cycle length by 43.8% (Fig. 3B, Fig. S6B). Of note, some FUCCI-WM164 cells displayed a delay in S/G2/M phase, which, however, did not reach statistical significance for the bulk population (Fig. 3B, Fig. S6C).

**Figure 3.**
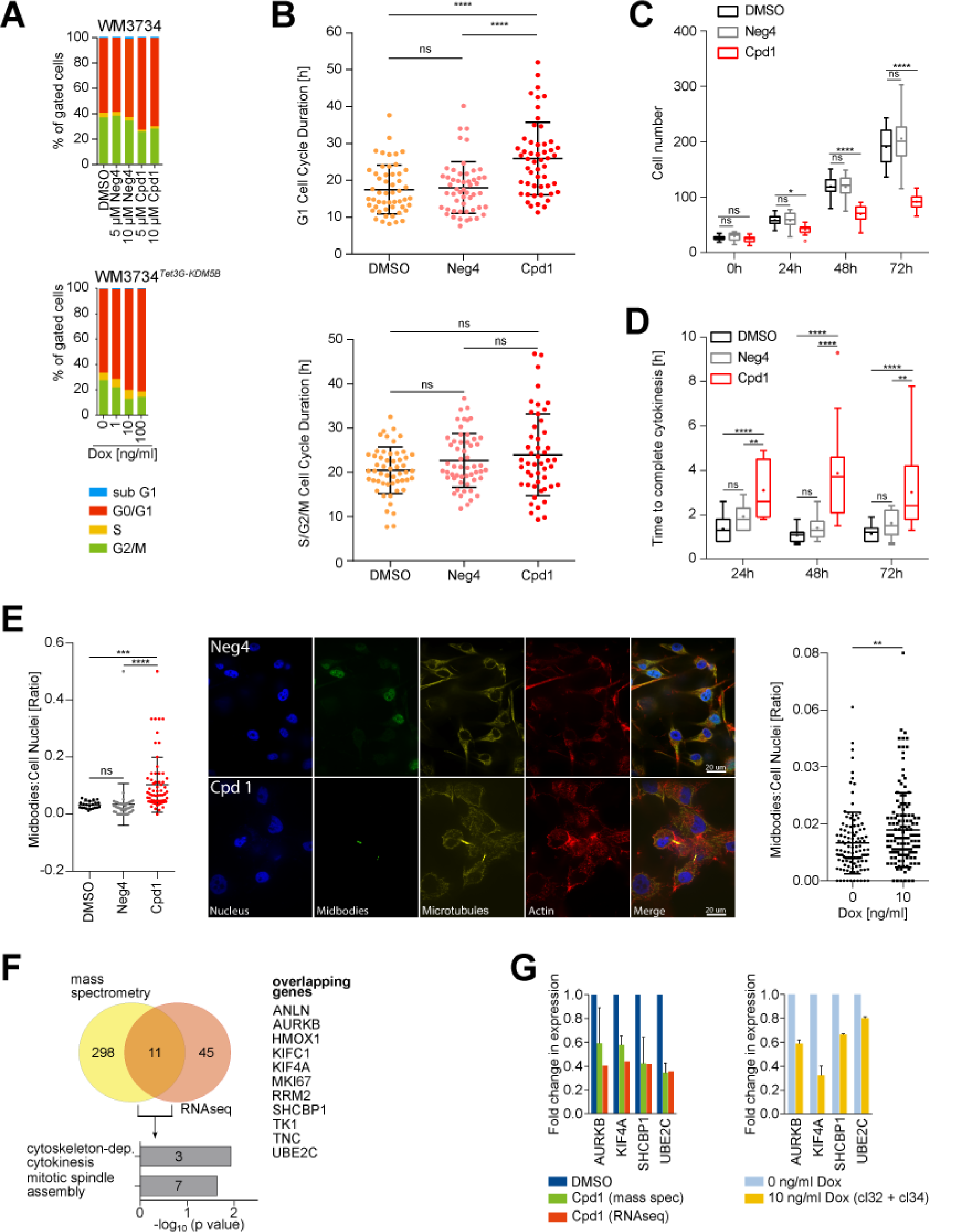
KDM5B-mediated cell cycle arrest and inhibition of cytokinetic abscission. (**A**) Propidium iodide flow cytometric cell cycle analysis of Cpd1-treated WM3734 (above) or Dox-treated WM3734*^Tet3G-KDM5B^* cells (below) after 72 h. Shown are representative data from 4 independent biological replicates. (**B**) Quantitation of the G1 (upper panel) and S/G2/M (lower panel) cell cycle duration by real-time cell cycle imaging of FUCCI-WM164 cells treated with Cpd1 (10 µM) vs. DMSO or Neg4 controls (10 µM) for 72 h. Scatter dot plots represent mean ±SD (50 cells of 5 replicates); ****P≤0.0001 by Kruskal-Wallis test. (**C**) Time-lapse imaging analysis of WM3734 cell numbers during treatment with Cpd1 (10 µM) vs. DMSO or Neg4 controls (10 µM) up to 72 h (15 different areas). Box-and-whiskers represent median values and the interquartile range; the mean values are plotted as asterisks; *P≤0.05, ****P≤0.0001 by Two-way ANOVA-test. This data reflect one experiment. (**D**) Time-lapse microscopic movies were analysed for the time to complete cytokinesis (15 different cells). Box-and-whiskers represent median values and the interquartile range; the mean values are plotted as <u>asterisks</u>; **P≤0.01, ****P≤0.0001 by Two-way ANOVA-test. (**E**) Quantitation and representative immunofluorescence staining of midbodies in Cpd1-treated WM164 cells (left) and Dox-treated WM3734*^Tet3G-KDM5B^* cells (right) vs. respective controls. Depicted are single fluorescence channels: blue (DAPI, nucleus), yellow (α-tubulin, microtubules); green (Aurora B kinase, midbodies); red (phalloidin, F-actin). Dot plots show the ratio of midbodies/cell per field of view, n= 12-16 fields from 3 independent replicates. Scatter dot plots represent mean ±SD; **P≤0.01, ***P≤0.001, ****P≤0.0001 by t-test. (**F**) Venn diagram and gene ontology analysis of significantly regulated genes upon Cpd1 (10 µM) treatment of WM3734 cells as detected by mass spectrometry and RNA sequencing. (**G**) Quantitation of downregulation of selected cytokinesis regulators as assessed by mass spectrometry and RNAseq (left) or qPCR (right) in WM3734 cells.

To further unravel the molecular mechanisms underlying the anti-proliferative effect of KDM5B, we performed single cell time-lapse microscopy of Cpd1-treated WM3734 cells vs. Neg4-treated control cells. The quantitation of cell numbers after 72 h confirmed a significant reduction in cell proliferation (Fig. 3C). Interestingly, we observed that some cells undergoing cell division, showed a prolonged time to complete cell abscission (Fig. 3D). We hypothesized that this phenomenon may be caused by a so far unknown regulatory role of KDM5B in cytokinesis (final process of physical separation of dividing cells). We aimed to quantitate this by immunostaining of intercellular midbodies, i.e. structures that represent microtubule-rich membrane bridges that connect two daughter cells shortly before their membranes fully dissever ^20^. Indeed, we found a significant increase in the number of midbodies in a fraction of Cpd1-treated WM164 cells (p<0.0001) as well as doxycycline-treated WM3734*^Tet3G-KDM5B^* cells (p=0.0033) (Fig. 3E). Transcriptional and proteomic profiling of KDM5B-enforced WM3734 cells by RNAseq and label-free quantitative mass spectrometry after 72 h of 10 µM Cpd1 revealed a small overlap of 11 genes/proteins of which 7 have known functions during cytokinesis (Fig. 3F, Table S2 highlighted in bold). For example, *AURKB*, *MKI67*, *KIF4A*, *UBE2C*, *RRM2*, *KIF1C1*, and *ANLN* are attributed to the GO cytoskeleton-dependent cytokinesis. For the expression of a subset of genes (*AURKB*, *KIF4A*, *SHCBP1*, and *UBE2C*), we found a downregulation in both Cpd1-treated WM3734 as well as doxycycline-treated WM3734*^Tet3G-KDM5B^* cells (Fig. 3G). *AURKB*, *KIF4A*, and *UBE2C* are known to regulate spindle assembly and coordinate abscission ^21, 22^. *SHCBP1* is involved in midbody organization and cytokinesis completion ^23^. Thus, irrespective of the model used, the above-mentioned genes may be functionally involved in the observed increase in midbodies and the delay in cell doubling.

Our experiments suggest so far that exogenous expression of KDM5B can direct melanoma towards a cell cycle-arrested tumor phenotype across the majority of tumor cells and several cell lines, where different molecular mechanisms might cooperate to prevent cell proliferation. Regarding potential clinical implications, enforcing the persister state could *per se* provide a therapeutic gain in time by delaying tumor progression. However, this approach might be more effective, if prolonged slow-cycling tumor cells are additionally eradicated by taking advantage of cell state-specific drug sensitivities.

### Enforced KDM5B expression leads to lineage reprograming

To unravel potential changes in cellular identity and underlying transcriptional programs after enforced KDM5B expression, we performed RNA sequencing of our genetic and chemical KDM5B induction models (Tet-On 3G and Cpd1) head-to-head. WM3734*^Tet3G-KDM5B^* cells were treated for 24, 48, and 72 h with doxycycline and then harvested for analysis. In parallel, naïve WM3734 and patient-derived short-term cultured CSM152 cells were treated with Cpd1 and analyzed after 72 h (Table S3). We first checked if Cpd1 also transcriptionally phenocopies the effect of Tet-On 3G-mediated KDM5B induction. Indeed, we found that both of our models can regulate a known KDM5B target gene motif to a similar degree ^24^ (Fig. S7A). Subsequent Gene Set Enrichment Analysis (GSEA) followed by Cytoscape visualization revealed a transcriptional landscape that matched other expected KDM5B-associated pathways like chromosomal remodeling or ATP-ase metabolism. Also genes involved in cell cycle/mitosis, DNA damage response, RNA processing, and immune response were found to be regulated by KDM5B (Fig. S7B). Transcripts that control cell cycle and mitosis revealed a steadily increasing and statistically significant regulation from 24 to 48 and 72 h of KDM5B induction pointing to a fundamental and temporally dynamic influence on the cell proliferation machinery (Fig. S7C).

As KDM5B has been reported to guide developmental processes in a highly conserved fashion ^25–, 27^, we asked if enforced KDM5B expression could affect the differentiation state of melanoma cells. Indeed, we observed for genetic as well as chemical KDM5B enforcement an impressive downregulation of mesenchymal and proliferative gene motifs (Fig. 4A, FWER p<0.05 was considered as statistically significant). Additionally, melanocytic differentiation genes were specifically regulated in both models (GO_PIGMENTATION, Fig. 4B). Most strikingly, enforced KDM5B expression was followed by a time-dependent transcriptional shift, in which cells were gradually transitioning from dedifferentiated to differentiated gene motifs over time (Fig. 4C, Table S4). Accordingly, KDM5B knockdown showed an inverse gene expression towards a dedifferentiated state (Fig. 4C). In an independent control experiment, differentiation reprograming was reproduced for Cpd1 and occurred also in a time-dependent manner (Fig. S7D, Table S4 and S5). Importantly, KDM5B-associated differentiation reprograming was similarly induced by Cpd1 in patient-derived CSM152 melanoma cells (Fig. 4A, B, Fig. S7A, S7D) and was also confirmed *in silico* for endogenous gene expression in single melanoma cells isolated from human tumor tissue (n=1253 melanoma cells from 19 tumors ^59^). Here, KDM5B expression was significantly higher in cells with melanocytic gene signatures compared to cells with undifferentiated signatures (Fig. 4D). As Cpd1 has been described in the literature as a calcium channel inhibitor, we performed GSEA of all model systems and excluded an overrepresentation of calcium and WNT-mediated signaling as off-target effects (KEGG_WNT_SIGNALING_PATHWAY, KEGG_CALCIUM_SIGNALING_ PATHWAY, GO_WNT_SIGNALING_ PATHWAY_CALCIUM_MODULATING_PATHWAY, data not shown).

**Figure 4.**
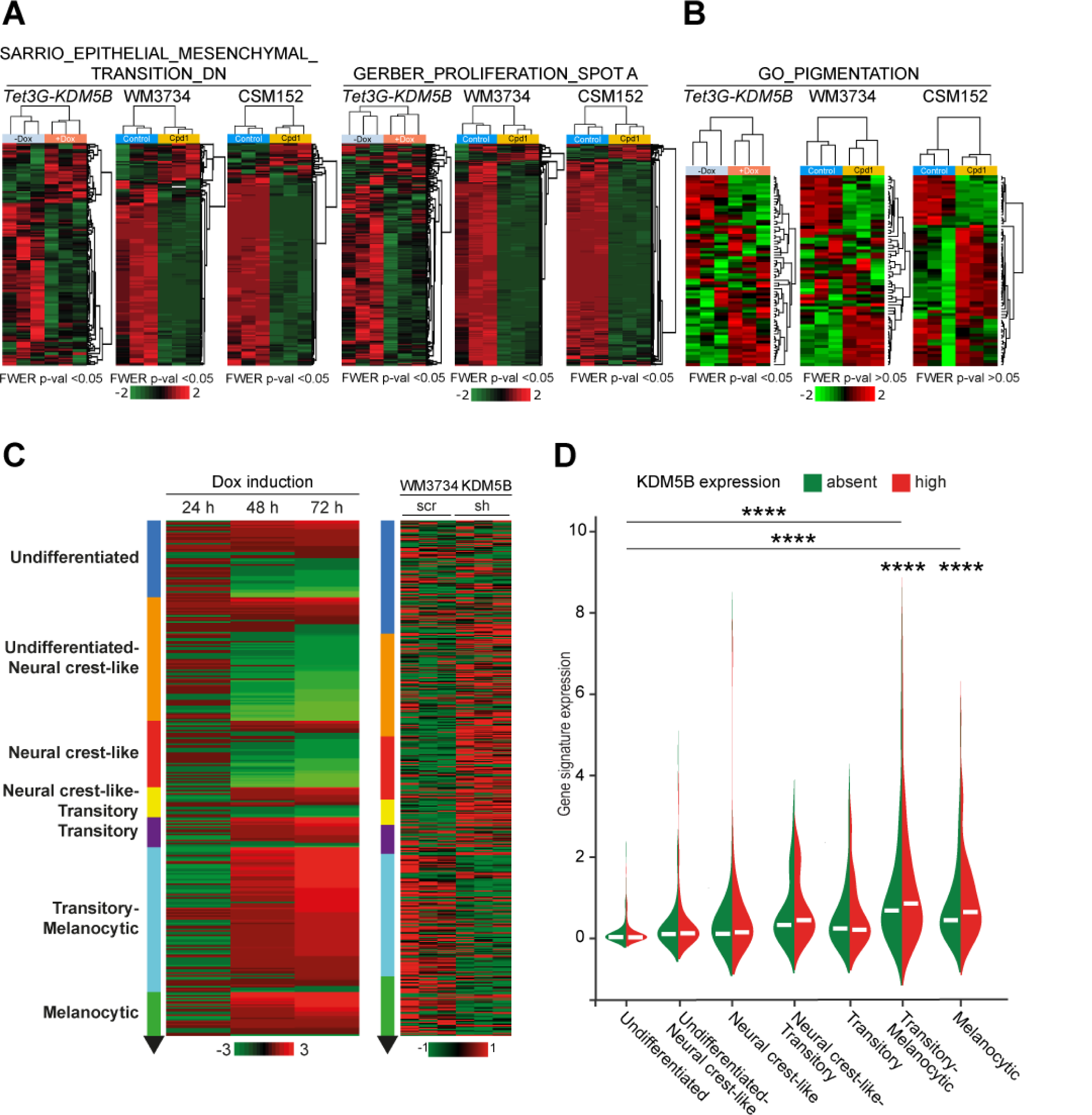
KDM5B leads to a differentiation-directed phenotypic shift. (**A** and **B**) Heatmaps of the SARRIO_EPITHELIAL_MESENCHYMAL_TRANSITION_DN ^47^, GERBER_PROLIFERATION_SPOT A signatures ^45^ (**A**) and GO_PIGMENTATION signature ^48, 49^ (**B**) from WM3734*^Tet3G-KDM5B^* cells after 72 h of Dox treatment and WM3734 and CSM152 cells after 72 h of Cpd1 treatment (red, upregulated; green downregulated genes). Significance is indicated by FWER P<0.05. (**C**) Heatmap of the Tsoi differentiation trajectory ^46^ for WM3734*^Tet3G-KDM5B^* cells after 24 h, 48 h and 72 h Dox treatment (left), WM3734 KDM5B knockdown (sh) versus control (scr) cells (right); red, upregulated, green downregulated genes. (**D**) Violin plot displaying endogenous KDM5B expression levels according to the transcriptional differentiation level in single cell RNA-sequenced human melanomas. Asterisks represent a p-value <0.0001 from 1-way Anova with Dunnett’s multiple comparison test. Here the comparison was made to the undifferentiated group.

### Enforced KDM5B expression leads to lineage-directed vulnerability

The observed KDM5B-dependent cell state reprograming not only confirms commitment to melanocytic differentiation, it could also sensitize melanoma cells for secondary phenotype-specific drugs. We sought to take advantage of 3-O-(3,4,5-trimethoxybenzoyl)-(-)-epicatechin (TMECG), a tyrosinase (TYR)-processed anti-metabolic agent previously described to eliminate melanoma cells in a lineage-specific way ^28^. Thus, we first assessed whether Cpd1 treatment truly activates the melanocytic differentiation/pigmentation machinery in melanoma cells. Indeed, we found a significant time-dependent Cpd1-induced transcription of the melanocytic master regulator MITF and other downstream differentiation genes like DCT, TYR, and MART-1/MLANA in pigmentation-competent melanoma cells (Fig. 5A). Conversely, knockdown of KDM5B was associated with a decrease in differentiation gene expression (Fig. 5B). Immunoblotting of WM3734*^Tet3G-KDM5B^* cells confirmed a KDM5B-dose-titratable increase of MITF protein, whereas typical markers of mesenchymal cell phenotypes such as N-cadherin, AXL, or ZEB1/2 were decreased (Fig. 5C). A similar regulation of these markers at protein level was confirmed for Cpd1 (Fig. 5D). Lastly, immunostained sections from syngeneic melanomas after Cpd1 treatment (Fig. 2L) confirmed enriched MITF protein expression *in vivo* (Fig. 5E, upper row). Fontana-Masson staining additionally indicated a focal increase of intracellular melanin production (Fig. 5E, lower row).

**Figure 5.**
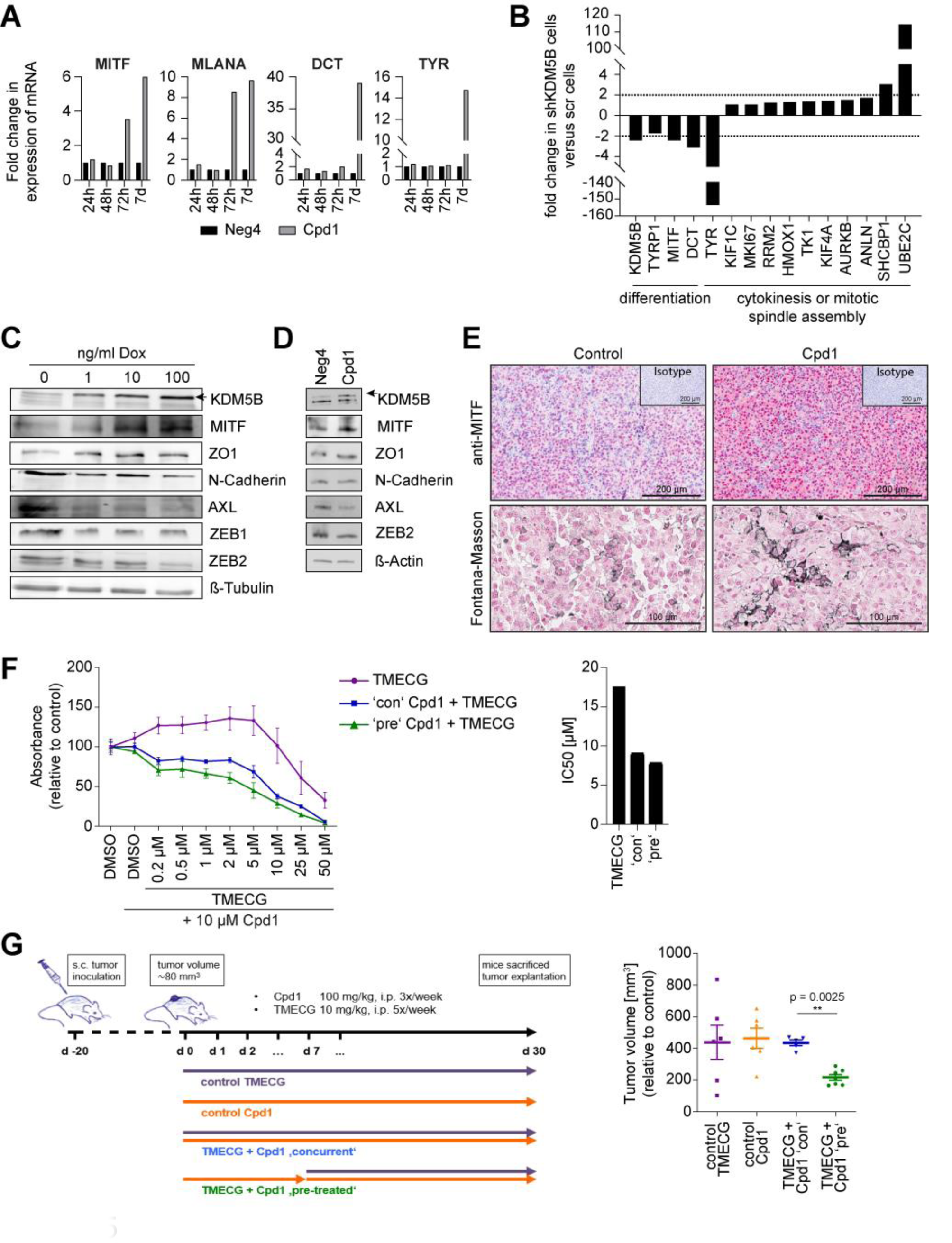
Enforced KDM5B expression facilitates melanocytic lineage-directed elimination by TMECG. (**A**) Quantitation of mRNA after 24 h, 48 h, 72 h and 7d of Cpd1 treatment of MaMel63a cells as assessed by qPCR. Mean ±SD. Shown is one representative example. (**B**) Regulation of differentiation, cytokinesis, and mitotic spindle assembly genes as detected by cDNA microarray analysis after KDM5B shRNA knockdown in WM3734 cells. (**C** and **D**) Immunoblotting of melanocytic lineage and (de-)differentiation markers after 24 h of KDM5B induction in WM3734*^Tet3G-KDM5B^* cells (**C**) and after 72 h of Cpd1 treatment in MaMel63a cells (**D**). Shown are representative data from two independent experiments. (**E**) Anti-MITF immunostaining (upper panel) and Fontana-Masson staining (lower panels) of CM melanoma tumor grafts from Cpd1-treated vs. control mice. (**F**) MTT cell viability assay of WM3734 cells. Representative example is shown left (n=2) and corresponding IC50 values on the right. TMECG was either concurrently given together with Cpd1 (‘con’) or added 3 days after Cpd1 pre-treatment (‘pre’). Readout was performed after 72 h of TMECG treatment. Data are shown as mean ±SD. (**G**) Persister-state-directed therapy model *in vivo*. Left: schematic representation of treatment dosing and timing in immunodeficient NMRI-(nu/nu)-nude mice. Right: tumor volumes of WM3734 xenografts (endpoint at day 30). TMECG was either concurrently given together with Cpd1 (‘con’) or added one week after Cpd1 pre-treatment (‘pre’). Data are shown as mean ±SEM. Significance was determined by Mann-Whitney test.

We next tested lineage-directed melanoma cell sensitization by Cpd1 in preclinical models. We tested combinations of Cpd1 with TMECG in three melanoma cell lines *in vitro* (WM3734, MaMel63a, WM983B, Fig. 5F and Fig. S8). MTT assays revealed that in the pharmacologically relevant, lower micromolar range of TMECG the combination with Cpd1 was particularly effective in cells, which were pre-treated with Cpd1 for 3 days. This compliments our *in vitro* observations that KDM5B-directed cell reprograming towards cell differentiation is a time-dependent process (Fig. 4C, Fig. S7D). TMECG alone showed a limited effect on cell numbers at low concentrations and, thus, was used as reference for subsequent *in vivo* experiments. We used an *in vivo* therapy model, in which WM3734 cells were xenografted to establish tumors on the back of immunodeficient NMRI-(nu/nu)-nude mice over 20 days (Fig. 5G left). Co-treatment was started either simultaneously for Cpd1 and TMECG or was performed consecutively, i.e. the tumors were pre-treated with Cpd1 for one week before TMECG was added. In line with our *in vitro* observations, melanoma tumor growth was significantly reduced when established tumors were primed to differentiation before state-specific elimination started (Fig. 5G right; p<0.05, Mann-Whitney test). Concurrent treatment of Cpd1 and TMECG failed to reduce tumor size.

To summarize, our study provides proof-of-concept for a new dual hit strategy in melanoma, in which persister state-directed transitioning limits cell state plasticity and primes tumor cells towards differentiation-specific elimination (Fig. 6).

**Figure 6.**
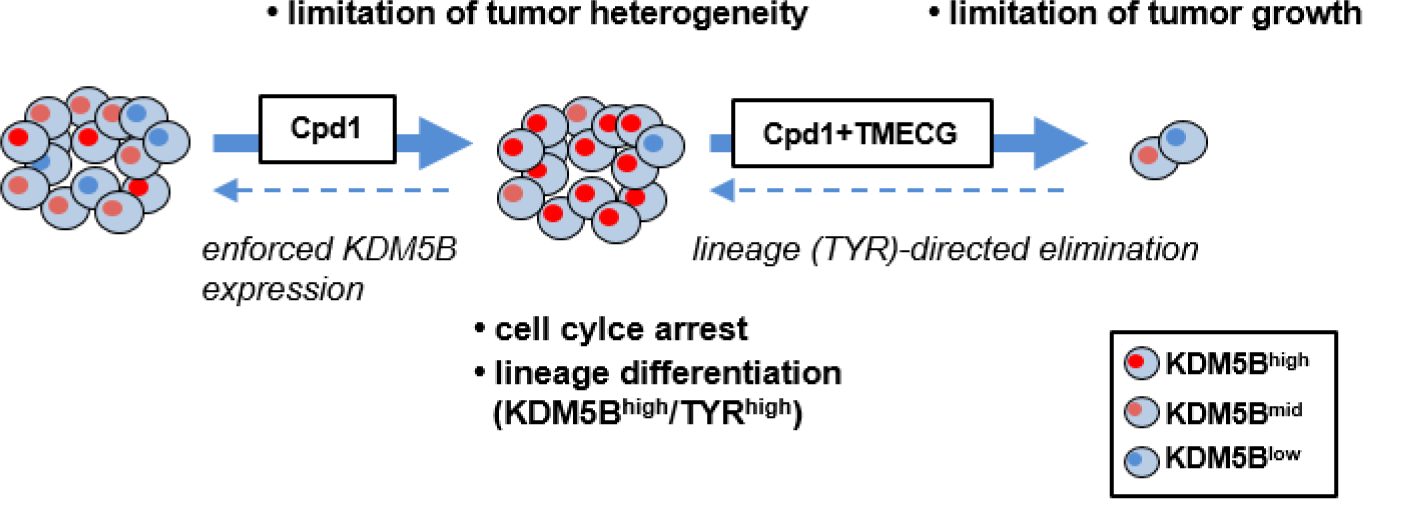
Phenotype-specific dual hit strategy in melanoma. KDM5B persister state-directed transitioning limits tumor plasticity and heterogeneity and primes melanoma cells towards lineage-specific elimination.

## Discussion

Next to genetic tumor evolution, particularly temporal phenotypic cell plasticity is an emerging problem in cancer therapy ^29^. As melanoma cells are rapidly shifting between different transcriptional programs, cell proliferation, and differentiation states, such phenotypes may transiently overlap and rapidly adapt to any strategy that specifically attacks single drug resistance mechanisms ^1–3, 30^. Thus, one question of this study was if limiting phenotypic cell state dynamics could become a starting point for a new dual hit strategy in melanoma. The slow-cycling KDM5B^high^ melanoma cell phenotype was used as a model for proof-of-concept.

So far, the real nature of slow-cycling KDM5B^high^ persisters in melanoma was unknown. Counter-intuitive to other currently discussed concepts of non-genetic tumor evolution such as transition to mesenchymal or stem-like states, our data strongly suggest that KDM5B favors transcriptional programs that drive a differentiated melanoma cell state similar to the recently reported DOT1L-inhibited persister state ^3^. In accordance with reports on KDM5B-dependent cell fate decisions and lineage commitment in other tissues types (e.g. neural differentiation ^26^ or hematopoiesis ^31^), our study suggests that KDM5B acts as a highly dynamic coordinator of both differentiation and cell division programs in melanoma. We show that KDM5B efficiently suppresses cell proliferation via downregulation of genes, which control cytoskeleton-dependent cytokinesis like *AURKB*, *KIF4A*, *UBE2C*, and *SHCBP1* ^21–23^. Our midbody analysis estimates that a considerable fraction of KDM5B-enforced melanoma cells have inhibited cytokinesis, which might contribute the observed G1 delay. As KDM5B is not affected by inactivating mutations in most melanomas according to currently available genetic profiling data like TCGA (mutation frequency of 5.9% according to CBioPortal), it may represent an ideal way for melanoma cells, irrespective of their mutational background to secure a slow-cycling state whenever required. Conceptually, this could mean that melanoma cells exploit the slow-cycling differentiated KDM5B^high^ state to immediately survive selective pressure, but must decrease KDM5B expression again and resume cell cycle progression in favor of more proliferative cell phenotypes to ensure long-term tumor repopulation (plasticity addiction).

Although we consider the KDM5B^high^ phenotype as differentiated, we are aware that any terminology that is applied to describe phenotypic cell states in melanoma needs to be used with caution because of their highly volatile nature. Under prolonged KDM5B induction, we observed changes in RNA signatures and protein marker profiles over time. Depending on the time point chosen for experimental read-out, marker constellations or even functional behavior like cell proliferation can be misleading for assessment of whether a certain cell state is advantageous or disadvantageous for tumor cell survival or drug killing. In this regard, our results unveiled a ‘Janus-faced’ role of the KDM5B^high^ melanoma cell state. On one hand, melanoma growth and cell invasion is impaired, when the slow-cycling state is sustained. On the other hand, melanoma cells can exploit this state to immediately survive targeted or cytotoxic therapies. However, as demonstrated here, when KDM5B^high^ state-directed transitioning is combined with a state-specific elimination strategy (instead of unspecific debulking drugs), control of tumor growth is possible. This represents a new alternative to current cytotoxic or signaling targeted therapies. In fact, multiple applications of cell state-directed elimination are conceivable now; particularly as a rescue strategy or to deepen response in residual disease after state-of-the-art cancer therapies. Thus, we plan to systematically test Cpd1/TMECG in sequential therapy regimens with other therapies in future studies ideally using a chemically improved version of Cpd1 with optimized solubility.

## Methods

### Melanoma cell lines, patient samples, and culture

The following human melanoma cell lines were maintained in 2% FBS-substituted (Tu2% ^32^) or 4% FBS-substituted (Tu4% ^33^ melanoma medium at 5% CO2: Wistar cell lines 451Lu (*BRAF^V^*^600^*^E^, PTEN^wt^, NRAS^wt^*), 451Lu BR (*BRAF^V600E^, PTEN^wt^, NRAS^wt^*), WM164 (*BRAF^V600E^, PTEN^wt^, NRAS^wt^*), WM3734 (*BRAF^V600E^, PTEN^del^, NRAS^wt^*), WM88 (*BRAF^V600E^, PTEN^wt^, NRAS^wt^*), WM9 (*BRAF^V600E^, PTEN^del^, NRAS^wt^*), WM983B (*BRAF^V600E^, PTEN^wt^, NRAS^wt^*), WM983B BR (*BRAF^V600E^, PTEN^wt^, NRAS^wt^*). Details on WM3734*^KDM5Bprom-EGFP^*, lentiviral infected WM3734_sh_KDM5B_62 and WM3734_sh_scramble control cells and WM3734^Tet3G-shJARID1B^ were previously described ^15, 34^. The commercial human melanoma cell lines MelJuSo (*BRAF^WT^, PTEN^WT^, NRAS^NRASQ61L^*), MeWo (*BRAF^WT^, PTEN^WT^, NRAS^wt^*), SKMel5 (*BRAF^V600E^, PTEN^n.d.^, NRAS^wt^*), SKMel28 (*BRAF^V600E^, PTEN^T167A^, NRAS^wt^*) were grown in RPMI medium with 10% FBS. The primary patient-derived melanoma cell lines CSM027 (*BRAF^V600E^, PTEN^wt^, NRAS^wt^*), CSM152 (*BRAF^wt^, PTEN^wt^, NRAS^wt^*), MaMel63a (*BRAF^V600E^, PTEN^wt^, NRAS^wt^*), ES014028 fibroblasts and the murine melanoma cell line CM ^18^ were also grown in RPMI medium with 10% FBS. Resistance of 451Lu BR and WM983B BR was maintained by 1 µM PLX4720 ^35^. Studies on human tissue samples and establishment of human melanoma cell lines were approved by the Internal Review Boards of the University of Pennsylvania School of Medicine and The Wistar Institute or the ethics committees of the Medical Faculties of the University of Wuerzburg and the University of Duisburg-Essen (reference numbers: 123/08_ff, 11-4715, 18-8301-BO). Cells were harvested using trypsin/EDTA 0.05%/0.02% in PBS (Biochrom). Cell line identity was confirmed by PCR-based DNA fingerprinting at the Department of Pathology of the University Hospital Essen. Cell culture supernatants were routinely tested for mycoplasma contamination using PCR with mycoplasma-specific primers. Analysis of human tumor samples was approved by the ethics committee of the Medical Faculty of the University of Duisburg-Essen (17-7373-BO).

### Establishment of an inducible Tet-On 3G-KDM5B cell line

A lentiviral Tet-On 3G-*KDM5B* construct was cloned for inducible KDM5B protein expression. Cloning steps were planned by VectorBuilder. The Tet-On transactivator protein is encoded by the *pLV-Hygro-CMV-Tet3G* vector. The TRE response vectors contain a PTRE3G promoter followed by either the human *KDM5B* gene, transcript variant 1, NM_001314042.1 (subcloned from *pBIND-RBP2-H1* ^36^ or, as control, the *EGFP* gene (*pLV-Puro-TRE3G-hKDM5B* and *pLV-Puro-TRE3G-EGFP*, respectively). In brief, WM3734 cells were stably infected with *pLV-Hygro-CMV-Tet3G* using JetPRIME Polyplus (NYC, NY, USA) according to the manufacture’s protocol. After selection by hygromycin B and single cell cloning, WM3734*^Tet3G^* cells were stably infected either with *pLV-Puro-TRE3G-hKDM5B* or *pLV-Puro-TRE3G-EGFP* followed by puromycin selection. Double-infected WM3734 melanoma cells (WM3734*^Tet3G-KDM5B^* or WM3734*^Tet3G-EGFP^*) were maintained in Tu2% media. Effects of doxycycline on the expression of MITF, the master regulator of melanocytic differentiation, were excluded by WB analysis (data not shown).

### Drugs and chemical compounds

The following drugs and compounds were used: ampicillin (AppliChem, Omaha, Nebraska, USA), blasticidin (InvivoGen, San Diego, CA, USA), cisplatin (1 mg/ml solution, Teva, Petach Tikwa, Israel), cycloheximide (Sigma Aldrich, St. Louis, Missouri, USA), DMSO (AppliChem, Omaha, Nebraska, USA), doxycycline (AppliChem, Omaha, Nebraska, USA), hygromycin (AppliChem, Omaha, Nebraska, USA), MG132 (Sigma Aldrich, St. Louis, Missouri, USA), PLX4720 (Selleckchem Houston, TX, USA), puromycin (Merck, Darmstadt, Germany), trametinib (Selleckchem Houston, TX, USA), Neg4 [(oxolan-2-yl)methyl 4-(6-bromo-2H-1,3-benzodioxol-5-yl)-2,7,7-trimethyl-5-oxo-1,4,5,6,7,8-hexahydroquinoline-3-carboxylate, ChemDiv, San Diego, CA, USA] and Cpd1 (2-phenoxyethyl 4-(2-fluorophenyl)-2,7,7-trimethyl-5-oxo-1,4,5,6,7,8-hexahydroquinoline-3-carboxylate, ChemDiv, San Diego, CA, USA). Neg4 and Cpd1 were dissolved in DMSO at a stock concentration of 10 mM and diluted 1:1,000 in media. 3-O-(3,4,5-trimethoxybenzoyl)-(-)-epicatechin (TMECG) was synthesized as described previously ^37^ and was made available by JN Rodríguez-López.

### Small chemical compound screening

To identify compounds which modulate expression levels of KDM5B in melanoma cells, a small molecule library was screened using our previously published KDM5B-promoter-*EGFP* reporter construct ^15^. The rationale was to find compounds, which decreased the reporter activity and, thus, the frequency of EGFP expressing cells below a threshold of 2% [K/EGFP, for threshold definition see also ^5, 15^]. We did not prioritize compounds that led to increased K/EGFP, since these were previously demonstrated to be possibly associated with unspecific cytotoxic effects ^5^. We were able to identify compounds, which yielded changes in transcriptional KDM5B levels upon compound treatment. The imaging screen was performed using an Opera (PerkinElmer, Waltham, Massachusetts, USA) High Content Screening system with confocal microplate imaging for readout and image analysis was performed using Columbus 2.4.0 (PerkinElmer, Waltham, Massachusetts, USA). The major measuring parameter of our assay was the K/EGFP level detected per cell and per well in relation to the total number of surviving cells after a 72 h treatment with compounds. Appropriate cell numbers used in the screen were 1,250 cells/well as determined by preliminary titration experiments with Draq5 identifying dead cells. Oligomycin (0.1 μg/ml) was used as a control for positive hits, i.e. compounds that decrease the fraction of K/EGFP expressing cells, trichostatin A (20 ng/ml) as negative control that absolutely increases K/EGFP without significant cell death, and cisplatin (20 μM) as negative control that relatively increases K/EGFP by killing bulk cells ^5^. The counter screen filtered out unspecific effects, i.e. only positive hits that were seen in WM3734*^KDM5Bprom-EGFP^* but not in WM3734*^CMVprom-EGFP^* control cells were considered specific.

The workflow comprised screening of 7,500 synthetic compounds from several well-known compound libraries, including the ENZO FDA approved drug library, (ENZO, 640 compounds), Analyticon Discovery library (AD, 2,329 compounds), ChemBioNet library (CBN, 2,816 compounds), ComGenex library (CGX, 2,437 compounds) and the Sigma-Aldrich Library of Pharmacologically Active Compounds (LOPAC, 1,280 compounds). The mechanism of action of some of these compounds are reported and known to modulate kinase, protease, ion channel and epigenetic regulators. 339 primary hits (AD: 46 compounds; CBN: 205 compounds; CGX: 2 compounds; ENZO: 18 compounds; LOPAC: 68 compounds) were identified, which decreased K/EGFP expressing cells < 2% without changing cell numbers more than 20% as compared to the DMSO control. In the next hit confirmation step, these 339 hits were confirmed in dose-response titration experiments in three independent runs with quadruplicates at each compound concentration (0.312, 0.625, 1.25, 2.5, 5.0 and 10 µM). The activities for 9 out of the 339 compounds met the criteria for progression, namely acceptable dose-response curve quality and potency. Out of the 9 validated hits, we chose one compound, 2-phenoxyethyl 4-(2-fluorophenyl)-2,7,7-trimethyl-5-oxo-1,4,5,6,7,8-hexahydroquinoline-3-carboxylate (termed Cpd1), to analyze the biological impact on melanoma cells. As a negative control for the biological assays we selected a structural analog compound (Neg4) that has a Molecular ACCess System (MACCS) similarity of 0.899, which was not identified as an hit compound (oxolan-2-yl)methyl-4-(6-bromo-2H-1,3-benzodioxol-5-yl)-2,7,7-trimethyl-5-oxo-1,4,5,6,7,8-hexahydroquinoline-3-carboxylate).

### Analysis of Cpd1

^1^H-NMR, ^13^C-NMR and ^19^F-NMR were recorded on *DRX600* (600MHz) spectrometer in MeOD. Data are reported in the following order: chemical shift (δ) in ppm; multiplicities are indicated s (singlet), d (doublet), t (triplet), q (quartet), m (multiplet); coupling constants (*J*) are given in Hertz (Hz). High resolution mass spectra were recorded on a LTQ Orbitrap mass spectrometer coupled to an Accela HPLC System (HPLC column: Hypersyl GOLD, 50 mm × 1 mm, 1.9 μm).

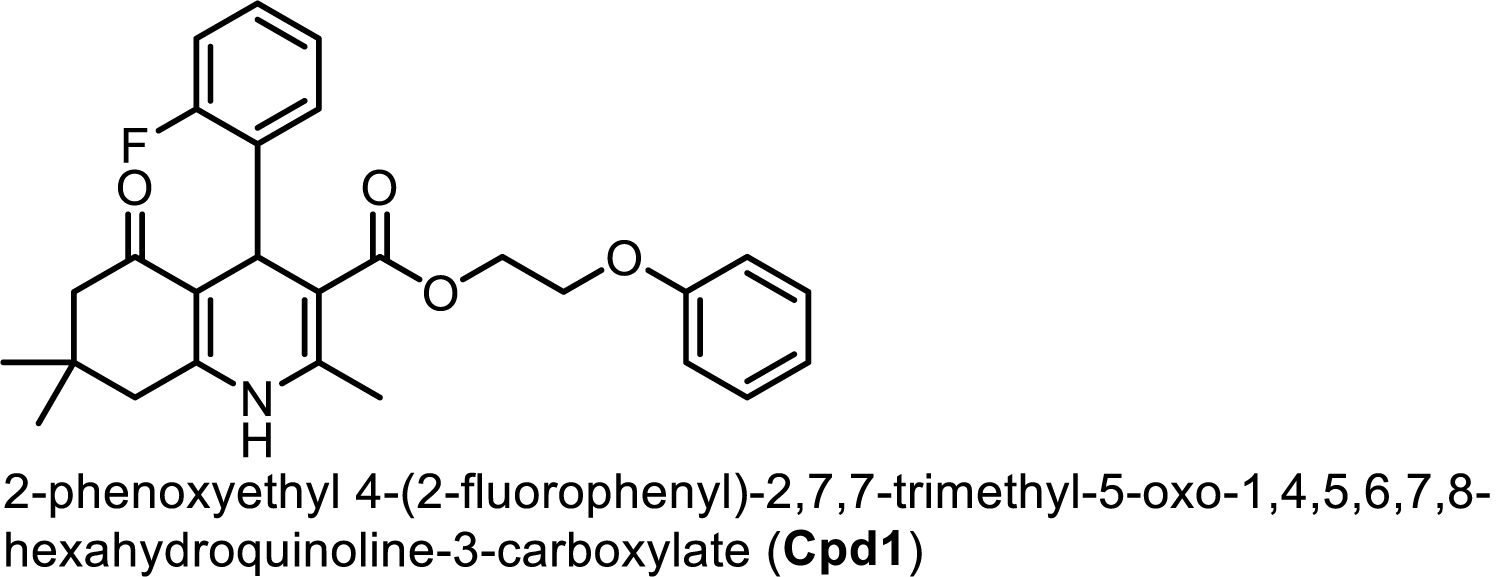

^1^H-NMR (600 MHz, MeOD) δ 7.29 (ddd, 1H), 7.02 (td, *J* = 7.3, 1.6 Hz, 1H), 6.94 (t, *J* = 7.3 Hz, 1H), 6.87 (dd, *J* = 16.9, 7.7 Hz, 1H), 6.79 (t, 1H), 5.18 (s, 1H), 4.34 (ddd, *J* = 12.0, 6.1, 3.3 Hz, 1H), 4.28 (ddd, *J* = 12.1, 6.2, 3.2 Hz, 1H), 4.14 (ddd, *J* = 9.5, 6.1, 3.2 Hz, 1H), 4.08 (ddd, 1H), 2.45 (d, *J* = 17.0 Hz, 1H), 2.32 (d, *J* = 15.0 Hz, 1H), 2.24 (d, *J* = 16.4 Hz, 1H), 2.04 (d, *J* = 16.4 Hz, 1H), 1.08 (s, 1H), 0.92 ppm (s, 1H). ^13^C-NMR (151 MHz, MeOD) δ 198.16, 169.01, 162.79, 161.15, 160.10, 152.99, 147.22, 135.03 (d, *J*_*CF*_ = 13.7 Hz), 132.59 (d, *JCF* = 4.8 Hz), 130.43, 128.76 (d, *J*_*CF*_ = 8.4 Hz), 124.47 (d, *JCF* = 3.3 Hz), 121.91, 116.14, 115.98, 115.64, 110.89, 104.76, 66.91, 63.44, 51.39, 41.02, 33.67, 33.39, 29.68, 26.87, 18.68 ppm. ^19^F-NMR (565 MHz, MeOD) δ -118.67 ppm. HR-MS: calc. for [M+H]^+^ C_27_H_29_O_4_NF = 450.20751 found 450.20785; C_27_H_28_O_4_NFNa = 472.18946 found 472.18962.

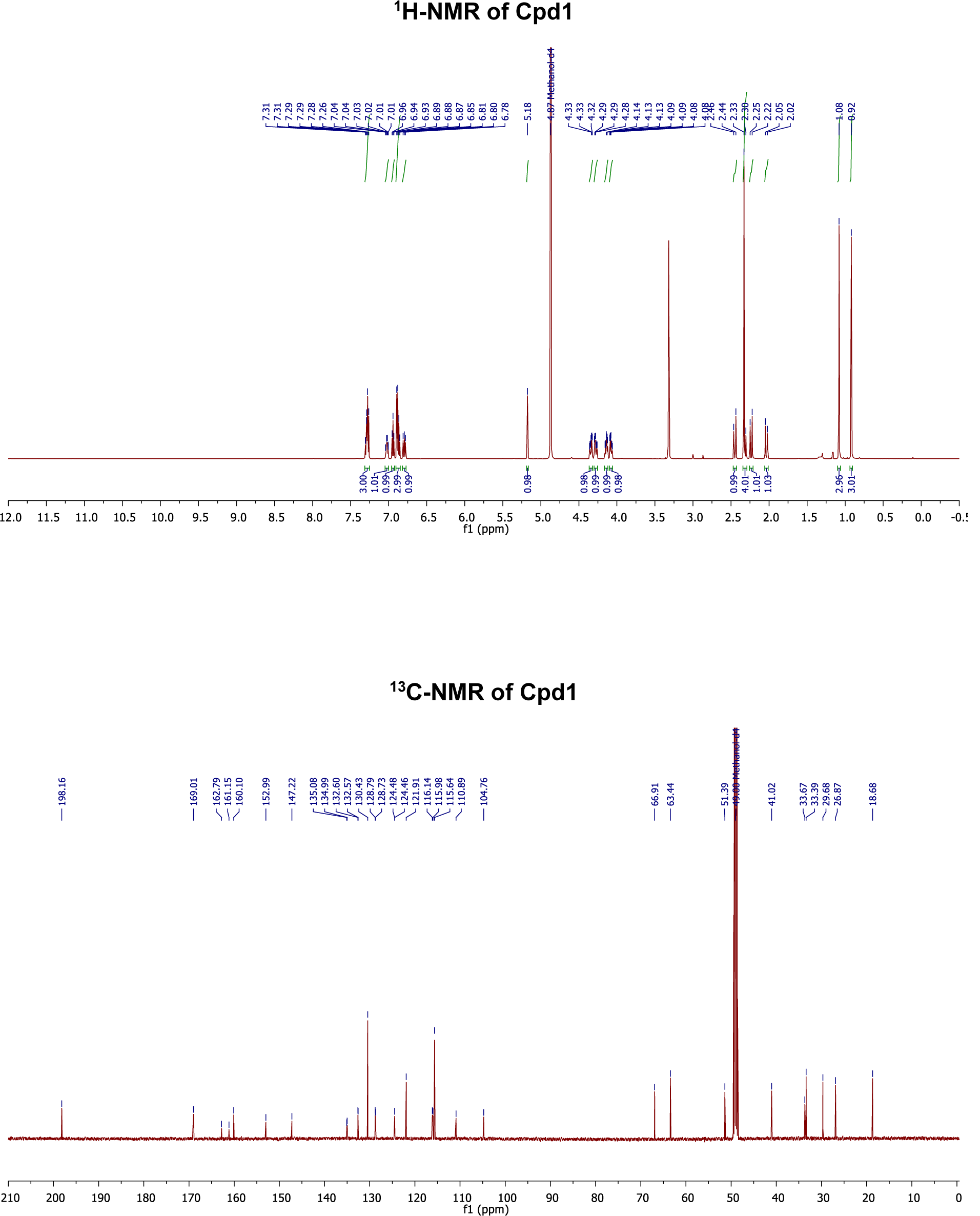

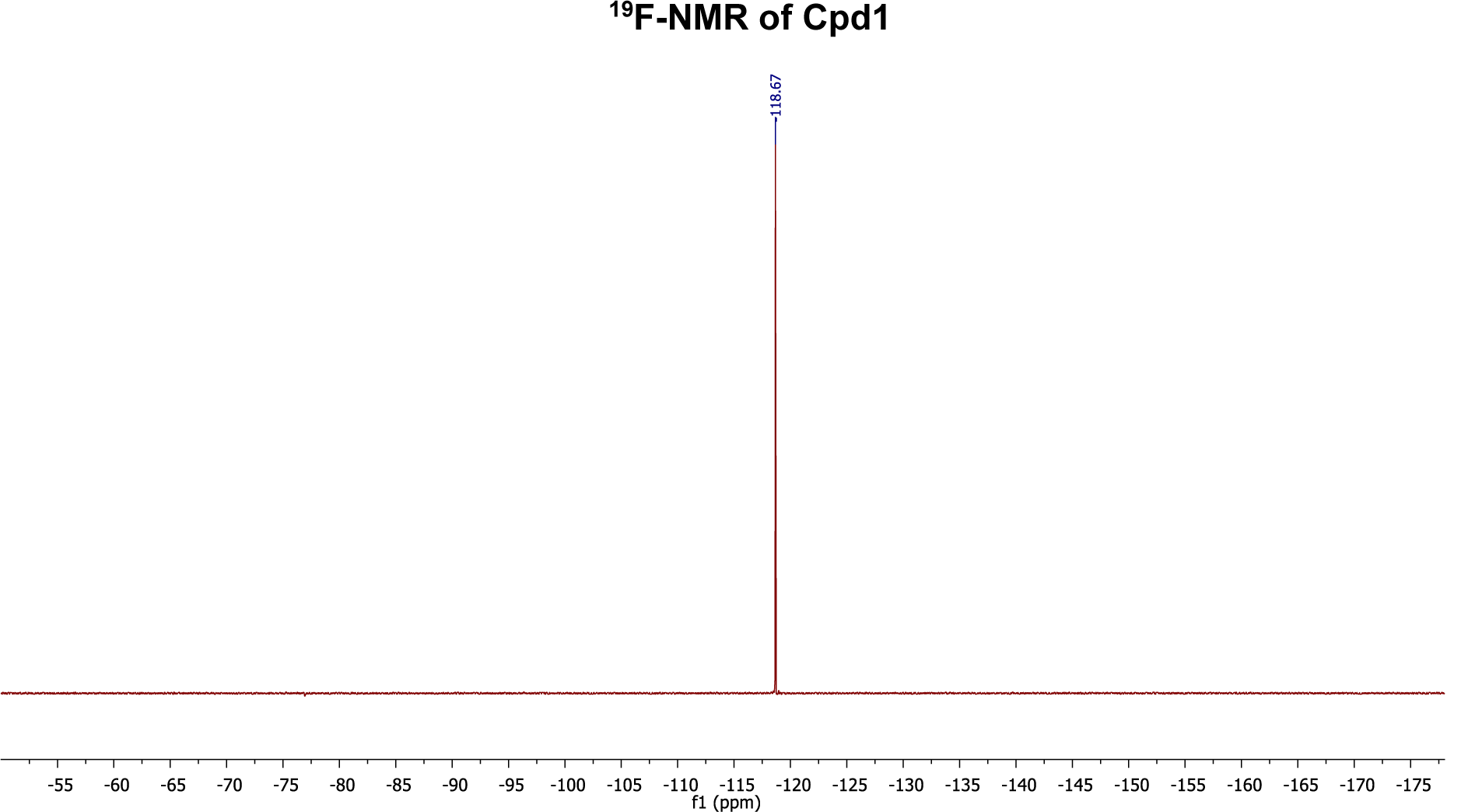

### Determination of cell numbers and apoptosis

Cell numbers were assessed by MTT and crystal violet assays according to standard protocols ^5^. Apoptosis was either detected by flow cytometry using LIVE/DEAD (ThermoFisher Scientific, Waltham, Massachusetts, USA) or IncuCyte apoptosis assays (caspase 3, Annexin V, ESSEN BioSCIENCE, Ann Arbor, Michigan, USA).

### Clonogenic, limited dilution, and colony formation assays

2D clonogenic growth was assessed after seeding 2,500 (used for 5-7-9 days readout) or 300 cells (used for 9-16-20 days readout) per 24-well plate. Cells were treated and analyzed by crystal violet (CV) staining. Quantitation of melanoma self-renewal was done by limited (single cell) dilution assays. In brief, cells were seeded at a density of 1 cell for every 4-wells in 96-well plates and grown for 19 days. Colony numbers were assessed microscopically by manual counting.

3D colony formation was assessed after 2,500 cells had been embedded into 0.35% soft agar in 6-well plates and grown over 30-52 days. Anchorage-dependent growth was inhibited by growing cells on a bed of 1% soft agar, with Tu2% or RPMI culture medium added on top and changed twice a week. For induction of KDM5B by doxycycline, WM3734*^Tet3G-KDM5B^* cells were pre-treated with doxycycline for 24 h before seeding and then continuously treated for the duration of the assay. For induction of KDM5B by the chemical modulator, melanoma cells were either continuously treated with the designated concentrations of Cpd1 in soft agar until colonies reached 3 mm in size (day 25) or pre-treated with 10 µM Cpd1 for 72 h before seeding in agar (pre-treatment). Colony numbers were assessed microscopically.

### Collagen-embedded melanoma spheroids and cell invasion assays

Melanoma spheroids were generated as described previously ^38^. In brief, 3,000 cells were grown in each well of a 96-well plate on top a layer of non-adherent 1.5% agarose for 72 h to form spheroids. The spheroids were then individually collected and embedded in a collagen type I mixture to assess 3D invasion. Spheroids were imaged 10 days after collagen embedding using a Zeiss AxioObserver.Z1 microscope. Spheroid invasion was quantitated using ImageJ 1.48x software (NIH). Normalized invasion was quantified by subtracting the spheroid volume area from the invasion area.

### Immunostaining of tissues

Paraffin embedded tumors or nevi were cut in 1.5, 2.5 or 4 µm sections. After deparaffinization and dehydration antibody staining was performed as indicated below or with silver nitrate working solution and fast red counterstain. Stained slides were scanned with an Aperio ScanScope AT2 (Leica) using 20x objectives and selected regions were selected in Aperio ImageScope software.

### Immunostaining of KDM5B and MITF protein

Cellular stainings were either performed by chemical or fluorescent staining. For chemical immunostaining of KDM5B, cells were seeded onto glass cover slips, fixed in 4% PFA with 0.1% Triton-X 100 for 10 min, and blocked with PBS containing 0.1% Triton-X 100 plus 5% BSA for 30 min. Staining was performed using anti-KDM5B NB100-97821 (Novus Biologicals, St. Louis, Missouri, USA, 1:1,200) and, as negative control, rabbit IgG (Dianova, Castelldefels, Spain, 1:1,200). The Dako REAL Detection System (Agilent, Santa Clara, CA, USA) was subsequently applied according to the manufacturer’s protocol and assessed using an Olympus BX51 or Zeiss AxioObserver.Z1 microscope. For immunostaining of KDM5B and MITF in FFPE tissue sections, slides were processed in a DAKO autostainer according to the manufacturer’s protocol using the same antibodies or anti-MITF ab12039 (Abcam, Cambridge, UK, 1:1,000). For immunofluorescence staining of KDM5B, cultured cells were fixed in 2% formalin containing 0.1% Triton-X 100 for 15 min at room temperature (RT) followed by blocking with 5% BSA containing 0.1% Triton-X 100 for 15 min at RT. Subsequent incubation with the primary antibody (1:1,200 to 1:20,000) or rabbit IgG (1:1,200 to 1:20,000, accordingly) was done for 1 h at RT. Alexa Fluor 568 goat anti-rabbit (1:600) served as secondary antibody. The staining was evaluated using a Zeiss AxioObserver.Z1 microscope. Image processing was applied equally across the entire image and was applied equally to controls.

### Immunofluorescence staining of midbodies

WM164 and WM3734*^Tet3G-KDM5B^* cells were grown in adherent sub-confluent culture and treated with Cpd1 (10 µM), DMSO, Neg4 control (10 µM) or 10 ng/ml doxycycline for 72 h, respectively. Samples were fixed with 4% paraformaldehyde in BRB80 buffer (80 mM K-PIPES, pH 6.8, 1 mM MgCl2, 1 mM EGTA) for 20 min, followed by three PBS washes and permeabilized with 0.25% Triton X-100 in PBS for 5 min. Samples were incubated in AbDIL (antibody dilution blocking buffer, 2% BSA, 0.1% Triton X-100, 0.1% NaN3 in PBS) overnight at 4°C. Primary antibodies to identify Aurora B kinase [1:100, (AIM1, 611082, BD Biosciences, Franklin Lakes, NJ, USA)], and α-Tubulin [1:400, (Abcam, Cambridge, UK)] and phalloidin [1:100, (Sigma-Aldrich, St. Louis, Missouri, USA)] were diluted in AbDIL and incubated overnight. Samples were washed three times prior to the addition of secondary antibodies diluted (1:500) in AbDIL containing 4 ′, 6-diamidine-2ʹ-24 phenylindole dihydrochloride [1:1,000 (Sigma-Aldrich, St. Louis, Missouri, USA)] for 1-2 h at RT. Samples were rinsed three times in PBS, 5 min each, and mounted in MOWIOL mounting medium (0.1 M Tris-HCl pH 8.5, 25% glycerol). Positive Aurora B Kinase and α-Tubulin stained midbodies were reported as a ratio of midbodies/cells per field of view, n = 12-16 fields. One field of view consists of 5 images in width x 5 images in length. Images were acquired using a custom-built Nikon Spinning Disk Confocal ^39^ or a Zeiss AxioObserver.Z1 microscope at 63x or 100x magnification (numerical aperture of Nikon Spinning Disk Confocal is 1.49, for AxioObserver.Z1 0.95 or 1.4). Image processing was applied equally across the entire image and was applied equally to controls.

### Quantitation of immunostaining

Immunofluorescence pictures were taken at 20x magnification using a Zeiss AxioObserver.Z1 microscope (numerical aperture is 0.8). To determine mean intensities of KDM5B signals nuclear and cytoplasmic KDM5B signals from at least 3 representative pictures for each independent experiment were quantitated using CellProfiler (Broad Institute Broad Institute, Cambridge, Massachusetts, USA) ^40^. Quantification of nuclear chromogen intensity of chemical immunostaining images was performed by using the reciprocal intensity method as published before ^41^.

### Flow cytometry

Flow cytometric analysis of KDM5B was done as described previously ^16^. Briefly, cells were harvested and fixed in 2% formalin with 0.1% Triton X-100 in PBS for 20 min at RT followed by permeabilization in 90% methanol for 30 minutes at –20°C. Primary [NB100-97821 (Novus Biologicals, St. Louis, Missouri, USA) or rabbit IgG (Dianova, Castelldefels, Spain)] and secondary antibodies [Alexa Fluor 568 goat anti-rabbit IgG or Alexa Fluor 647 goat anti-rabbit IgG (Life Technologies, CA, USA)] were incubated for 30 min at RT. Before and after antibody incubation, 25 cells were washed with FACS buffer (PBS containing 0.5 M EDTA and 1% FBS). Samples were measured utilizing a Gallios flow cytometer (Beckman Coulter, Brea, CA, USA) and analyzed with Kaluza 1.2 (Beckman Coulter, Brea, CA, USA) or FlowJo V7.6.5 (Tree Star, Ashland, Oregon, USA) software. Gates were set based on the DMSO control, which was set to 1% KDM5B^high^ cells. For flow cytometric detection of KDM5B promoter-driven *EGFP* signals, WM3734*^KDM5Bprom-EGPF^* cells were harvested, washed with FACS buffer, and stained for 7-AAD (eBioscience, San Diego, CA, USA). For quantitation, a 5%-threshold for the K/EGFP signal intensity was applied as described previously ^5, 15^.

### Cell cycle profiling

Cell cycle analysis was performed by propidium iodide staining as described previously ^15^. In brief, 100,000 MelJuSo, WM9, or SKMel5 cells were seeded per T75 flask or 20,000 WM3734 cells per 6 cm dish and starved for 5 days in medium without FBS. Starting from day 6, cells were treated for 72 h with Cpd1 either in the presence or absence of 2% FBS. WM3734*^Tet3G-KDM5B^* clones (80,000 cells) were seeded per 6 cm dish and starved for 5 days. Starting from day 6, doxycycline (0, 1, 10 or 100 ng/ml) was added in FBS containing medium and replaced every 2 to 3 days. Cells were analyzed after 6 days of treatment. For propidium iodide staining, cells were trypsinised and washed with PBS containing 5 mM EDTA. Cells were then fixed with 100% ethanol for 30 min at RT followed by a RNase A treatment for 30 min at RT. Propidium iodide was added at a final concentration of 100 µg/ml. Quantitation was done on a Gallios cytometer (Beckman Coulter, Brea, CA, USA) in linear mode. Data analysis was performed using FlowJo V7.6.5 or Kaluza 1.2 software.

### Fluorescent ubiquitination-based cell cycle indicator (FUCCI)

To generate stable melanoma cell lines expressing the FUCCI constructs, *mKO2-hCdt1* (30-120) and *mAG-hGem* (1-110) ^42^ were sub-cloned into a replication-defective, self-inactivating lentiviral expression vector system as previously described ^32^. The lentivirus was produced by co-transfection of human embryonic kidney 293T cells. High-titer viral solutions for *mKO2-hCdt1* (30/120) and *mAG-hGem* (1/110) were prepared and used for co-transduction into melanoma cell line WM164 and sub-clones were generated by single cell sorting ^19, 33^.

### Time-lapse microscopy

To track cell numbers, 80,000 WM3734 cells were seeded per 6 cm dish and images were taken every 5 minutes over 5 days on 15 different areas per condition at 10x magnification using a Zeiss AxioObserver.Z1 microscope. Cell numbers were manually counted for every position after 0, 24, 48 and 72 h of treatment. Cell division time was determined by measuring the time (in min) between cell rounding and complete abscission of daughter cells.

FUCCI-WM164 cells ^19^ were grown in 24-well dishes and pre-treated with Cpd1 (10µM), Neg4, or DMSO for 24 h. Images for time lapse were acquired every 10 min over 72 h from 5 different positions in triplicate wells per experiment. FUCCI probe fluorescence intensity over time were calculated as previously published ^43^ with the following modifications. Total duration and profile of FUCCI fluorescence of the different stages of the cell cycle were determined using the ‘spots function‘ in IMARIS, Bitplane. A threshold for spots equal-to or less-than 15 µm were used to track fluorescent FUCCI nuclei utilizing the auto aggressive motion algorithm. Track paths, following fluorescent nuclei, were then used to create ROIs and measure mean center point intensities throughout the duration of the movie. Tracks greater than 50% of the total movie duration were used for analysis to ensure full cell cycle profiles were captured. Tracks with breaks greater than 2 frames were disregarded and cells coinciding within 30 µm of peripheral X-Y borders were ignored to avoid including partial track profiles. Mean fluorescence intensity values were background fluorescence corrected and used to calculate total track duration and generate plots of intensity variation over time.

### Real time quantitative RT-PCR (QPCR)

Total RNA was isolated using the RNeasy Mini Kit according to the manufacturer’s protocol (Qiagen, Venlo, Netherlands). 20 ng RNA was used as template for quantitative PCR with Precision OneStep qRT-PCR master mix (PrimerDesign, Southampton, UK). Quantitative PCR was performed with a StepOnePlus Real-Time PCR system (ThermoFisher Scientific, Waltham, Massachusetts, USA). Thermal cycler conditions were 95°C for 20 min, then 40 cycles of 3 min at 95°C, followed by 30 sec at 60°C. The analysis was performed using the StepOnePlus software (ThermoFisher Scientific, Waltham, Massachusetts, USA). mRNA expression was calculated using the 2^-DDCT^ method and normalized to the housekeeping control 18S. Used primers are listed in Table S6.

### Immunoblotting

For immunoblotting of whole cell lysates, cells were either lysed with RIPA buffer (50 mM Tris-HCl, pH 6.8, 150 mM NaCl, 0.5% sodium deoxycholate, 0.1% SDS and 1% Triton X-100) supplemented with phosphatase inhibitors (cOmplete tablets, Roche Diagnostics) or according to the REAP protocol ^44^. Samples (20-25 µg of protein) were separated on 8% polyacrylamide-SDS, wet transferred onto PVDF membranes (Roth, Karlsruhe, Germany) and blocked for 1 h in 5% milk containing 0.1% Tween-20. Primary antibodies [N-cadherin (13116), phospho-Rb (9309), phospho-Rb (Ser780; 8180), phospho-Rb (Ser795; 9301), phospho-Rb (Ser807/811; 8516), tubulin (2148), Rb (9309), ZEB1/TCF8 (3396), ZEB2/Smad1 (9743) and ZO-1 (8193; all diluted 1:1,000, all Cell Signaling, Cambridge, UK), histone H3 (ab1791, diluted 1:5,000), histone H3K4me3 (ab8580, diluted 1:2,000), MITF (ab80651, diluted 1:500) and Notch1 (ab52627, all diluted 1:1,000, all Abcam, Cambridge, UK), KDM5B (NB100-97821, diluted 1:2,000, Novus Biologicals, St. Louis, Missouri, USA) and GAPDH (SC-510, diluted 1:5,000, Santa Cruz, Dallas, Texas, USA)] were incubated overnight at 4°C either in PBS containing 0.1% Tween-20 and 5% milk or in 1x Net-G buffer (10x Net-G contains 1.5 M NaCl, 50 mM EDTA, 500 mM Tris 0.5% Tween 20 and 0.4% gelatine). Blots were washed with PBS-T or NetG followed by a 1 h incubation with horseradish peroxidase-conjugated secondary antibody [anti-rabbit (115-035-046) or anti-mouse (115-035-003, Jackson Immuno Research Laboratories, West Grove, PA, USA)] diluted 1:10,000 in PBS-T-milk 5% or NetG and a further washing step. Bands in Western blots were visualized by an enhanced chemiluminescence system (WesternBright Chemiluminescence Substrate, Advansta, Menlo Park, CA, USA) and captured using a FUJI LAS3000 system. Digital quantitation was performed using ImageJ 1.48x software (NIH).

### In vivo studies

All animal experiments were performed in accordance with institutional and national guidelines and regulations. The protocols have been approved by the local German authority Landesamt für Natur, Umwelt und Verbraucherschutz Nordrhein-Westfalen – LANUV NRW in compliance with the German animal protection law (Reference number AZ 84-02.04.2014.A08 AZ 81-02.04.2018.A202). The maximum tolerable dose of Cpd1 was determined in a prior experiment. Here, mice (n=5) did not show abnormal behavior or body weight loss up to 100 mg/kg/d. Xenograft tumors of the human melanoma cell lines WM3734 or WM3734^Tet3G-KDM5B^ were generated by injection of 2x10^5^ cells in 200 µl medium [1:1 mixture of Tu2% with Matrigel® (BD Biosciences, Franklin Lakes, NJ, USA)] s.c. on the back of immunodeficient NMRI-(nu/nu)-nude mice. WM3734^Tet3G-KDM5B^ xenograft model: once tumors reached 150 mm^3^ by caliper measurement (calculated as WxWxL/2), animals were randomized into 2 groups, ‘control’, and ‘500 Dox’, and the drinking water was supplemented accordingly with 2.5% sucrose plus 500 µg/ ml doxyclycline (provided *ad libitum*). Control mice received 2.5% sucrose-substituted water without doxycycline. Water was changed twice a week. WM3734 xenograft model: when tumors became palpable, animals were randomized into 4 groups, ‘control Cpd1’ (100 mg/kg 3x/week (i.p.), ‘control TMECG’ (10 mg/kg 5x/week (i.p.), ‘TMECG plus Cpd1 con’ (10 mg/kg 5x/week and 100 mg/kg 3x/week, respectively) and ‘TMECG plus Cpd1 pre’ (10 mg/kg 5x/week starting 7 days later and 100 mg/kg 3x/week, respectively). Tumor growth was measured three times a week using a caliper. Tumor samples were fixed in formalin for histological assessment and immunostaining. Xenograft tumors of the murine melanoma cell line CM ^18^ were generated by injecting 1x10^5^ cells in 200 µl medium [1:1 mixture of RPMI medium with Matrigel® (BD Biosciences, Franklin Lakes, NJ, USA)] s.c. on the back of female C57BL/6N mice. Once tumors reached 400 mm^3^ by caliper measurement (calculated as WxWxL/2), animals were randomized into 2 groups with each 5 mice, ‘Cpd1’ (100 mg/kg) every second day intraperitoneal (i.p.), and ‘control group’ (PEG300+IgG, 250 µg every second day i.p.). Dosing continued until tumors had reached the maximal volume. Tumor samples were fixed in formalin for histological assessment and immunostaining.

### RNAseq transcriptional profiling

For generating total RNA either WM3734 and CSM152 (100,000 cells) were seeded in a 6 cm dish and treated with Cpd1 (10µM) or DMSO for 12 h, 24 h, 48 h, or 72 h. WM3734*^Tet3G-KDM5B^* (40,000 cells) were seeded in a 6 cm dish and induced with 10 ng/ml doxycycline for 24 h, 48 h, or 72 h. WM3734^Tet3G-shJARID1B^ or as control WM3734^Tet3G-scramble^ (175,000 cells) were seeded in a 6 cm dish and induced with 500 ng/ml doxycycline for 72 h. Total RNA was isolated using RNeasy Mini Kit according to the manufacturer’s protocol (Qiagen Venlo, Netherlands). Barcoded stranded mRNA-seq libraries were prepared using the Illumina TruSeq RNA Sample Preparation v2 Kit (Illumina, San Diego, CA, USA) implemented on the liquid handling robot Beckman FXP2. Obtained libraries were pooled in equimolar amounts; 1.8 pM solution of this pool was loaded on the Illumina sequencer NextSeq 500 and sequenced uni-directionally, generating 500 million reads 85 bases long. The run was basecalled and demultiplexed using Illumina bcl2fastq2, version 2.20.0.422. The alignment was done using BWA mem, version 0.7.17. with default parameters. The reference genome for alignment was hg19 or in case of WM3734^Tet3G-shJARID1B^ cells hg38. Finally, statistical gene set analysis was performed to determine differential expression at both gene and transcript levels. Partek flow defaults were used in all analyses. Gene set enrichment analysis (GSEA-http://software.broadinstitute.org/gsea/index.jsp) was performed using the pre-ranked tool (Broad Institute, Cambridge, Massachusetts, USA). All genes which contained an average of more than 1 read across all samples were used and ranked according to the T statistic. Gene sets were comprised of curated pathways from several databases including GO, Reactome, KEGG (March 24 2016 version; http://download.baderlab.org/EM_Genesets/current_release/Human/symbol/) and visualized using Cytoscape (www.cytoscape.org; p<0.003, q<0.04, similarity cutoff 0.5). RNAseq of the cell lines WM3734 and CSM152 were analyzed separately and analyses were merged keeping only overlapping networks. Heatmaps were generated in Partek Genomic Suite or R using previously published genes sets ^24, 45–49^. Hierarchical clustering was performed by normalizing mean expression to 0 with a standard deviation of 1 and using Pearson’s dissimilarity algorithm and average linkage. The violin plot was generated in BioVinci.

### Transcriptional profiling by microarray

RNA from melanoma cells (WM3734 stably infected with shKDM5B (62) or as control scramble (SCR) ^5^ was extracted with Trizol reagent, followed by clean-up and DNase I treatment with QIAGEN RNeasy mini kit in accordance with the prescribed protocol provided with the kit. Micorarray transcriptional analysis was performed using the HumanWG-6 v3.0 expression BeadChip sytem (Illumina, San Diego, CA, USA) at the Wistar Genomics facility. The data were processed with Illumina GenomeStudio Gene Expression Module using defaults.

### Sample preparation and clean-up for LC-MS

For LC/MS proteomic analysis, 400,000 WM3734 cells were seeded per 10 cm dish and treated on day 4 with Cpd1 (10 µM) or DMSO for 72 h. Cells were washed with PBS once before harvesting by mechanical detachment in PBS. Cells were centrifuged at 2,000 rpm for 4 min and the subsequent pellet resuspended in 120 µl lysis buffer [RIPA buffer (50 mM Tris-HCl, pH 6.8, 150 mM NaCl, 0.5% sodium deoxycholate, 0.1% SDS and 1% Triton X-100) supplemented with phosphatase inhibitors (cOmplete tablets, Roche Diagnostics)]. After 30 min incubation on ice, lysates were centrifuged at 13,000 rpm for 30 min and the supernatant was stored at –80°C. The samples were next reduced with DTT and alkylated with iodoacetamide and subsequently digested in the presence of sequencing grade LysC (Wako) and Trypsin (Promega, Fitchburg, WI, USA). Finally the acidified tryptic digests were desalted on home-made 2 disc C18 StageTips as described ^50^. After elution from the StageTips, samples were dried using a vacuum concentrator (Eppendorf, Hamburg, Germany) and the peptides were taken up in 10 µl 0.1% formic acid solution.

### LC-MS/MS settings

Experiments were performed on an Orbitrap Elite instrument (ThermoFisher Scientific, Waltham, Massachusetts, USA) ^51^ that was coupled to an EASY-nLC 1000 liquid chromatography (LC) system (ThermoFisher Scientific, Waltham, Massachusetts, USA). The LC was operated in the one-column mode. The analytical column was a fused silica capillary (75 µm × 30 cm) with an integrated PicoFrit emitter (New Objective) packed in-house with Reprosil-Pur 120 C18-AQ 1.9 µm resin (Dr. Maisch). The analytical column was encased by a column oven (Sonation) and attached to a nanospray flex ion source (ThermoFisher Scientific, Waltham, Massachusetts, USA). The column oven temperature was adjusted to 45°C during data acquisition. The LC was equipped with two mobile phases: solvent A (0.1% formic acid, FA, in water) and solvent B (0.1% FA in acetonitrile, ACN). All solvents were of UPLC grade (Sigma-Aldrich, St. Louis, Missouri, USA). Peptides were directly loaded onto the analytical column with a maximum flow rate that would not exceed the set pressure limit of 980 bar (usually around 0.6–1.0 µl/min). Peptides were subsequently separated on the analytical column by running a 140 min gradient of solvent A and solvent B (start with 7% B; gradient 7% to 35% B for 120 min; gradient 35% to 100% B for 10 min and 100% B for 10 min) at a flow rate of 300 nl/min. The mass spectrometer was operated using Xcalibur software (version 2.2 SP1.48). The mass spectrometer was set in the positive ion mode. Precursor ion scanning was performed in the Orbitrap analyzer (FTMS; Fourier Transform Mass Spectrometry) in the scan range of m/z 300-1800 and at a resolution of 60,000 with the internal lock mass option turned on [lock mass was 445.120025 m/z, polysiloxane; ^52^]. Product ion spectra were recorded in a data dependent fashion in the ion trap (ITMS) in a variable scan range and at a rapid scan rate. The ionization potential (spray voltage) was set to 1.8 kV. Peptides were analyzed using a repeating cycle consisting of a full precursor ion scan (3.0 × 10^6^ ions or 50 ms) followed by 15 product ion scans (1.0 × 10^4^ ions or 50 ms) where peptides are isolated based on their intensity in the full survey scan (threshold of 500 counts) for tandem mass spectrum (MS2) generation that permits peptide sequencing and identification. Collision induced dissociation (CID) energy was set to 35% for the generation of MS2 spectra. During MS2 data acquisition dynamic ion exclusion was set to 120 sec with a maximum list of excluded ions consisting of 500 members and a repeat count of one. Ion injection time prediction, preview mode for the FTMS, monoisotopic precursor selection and charge state screening were enabled. Only charge states higher than 1 were considered for fragmentation.

### Peptide and Protein Identification using MaxQuant

RAW spectra were submitted to an Andromeda ^53^ search in MaxQuant (1.5.3.30) using the default settings ^54^. Label-free quantitation and match-between-runs was activated ^55^. The MS/MS spectra data were searched against the Uniprot human reference database (UP000005640_9606.fasta, 70244 entries). All searches included a contaminants database search (as implemented in MaxQuant, 245 entries). The contaminants database contains known MS contaminants and was included to estimate the level of contamination. Andromeda searches allowed oxidation of methionine residues (16 Da) and acetylation of the protein N-terminus (42 Da) as dynamic modifications and the static modification of cysteine (57 Da, alkylation with iodoacetamide). Enzyme specificity was set to “Trypsin/P” with two missed cleavages allowed. The instrument type in Andromeda searches was set to Orbitrap and the precursor mass tolerance was set to ±20 ppm (first search) and ±4.5 ppm (main search). The MS/MS match tolerance was set to ±0.5 Da. The peptide spectrum match FDR and the protein FDR were set to 0.01 (based on target-decoy approach). Minimum peptide length was 7 amino acids. For protein quantitation unique and razor peptides were allowed. Modified peptides were allowed for quantitation. The minimum score for modified peptides was 40. Label-free protein quantitation was switched on, and unique and razor peptides were considered for quantitation with a minimum ratio count of 2. Retention times were recalibrated based on the built-in nonlinear time-rescaling algorithm. MS/MS identifications were transferred between LC-MS/MS runs with the “match between runs” option in which the maximal match time window was set to 0.7 min and the alignment time window set to 20 min. The quantitation is based on the “value at maximum” of the extracted ion current. At least two quantitation events were required for a quantifiable protein. Further analysis and filtering of the results was done in Perseus v1.5.5.1. ^56^. For quantitation we combined related biological replicates to categorical groups and investigated only those proteins that were found in at least one categorical group in a minimum of 5 out of 6 biological replicates. Comparison of protein group quantities (relative quantitation) between different MS runs is based solely on the LFQ’s as calculated by MaxQuant (MaxLFQ algorithm; ^55^.

### Statistical analysis

Overall survival curves were calculated from the TCGA data set (http://cancergenome.nih.gov/) using the TCGA browser tool v.0.9 (beta-testing version, http://tcgabrowser.ethz.ch:3839/TEST/). Unpaired t-tests were used to compare mean differences between groups. One-way ANOVA or two-way ANOVA was used to evaluate the association between different treatment groups. The unpaired t-test, one-way and two-way ANOVA test were performed in GraphPad Prism (version 6.07), conducted at the two-sided significance level, where p-values of <0.05 were considered significant. The tumor volumes of mice measured over time were used to reflect the tumor growth trend affected by different treatments. The velocities of tumor growth were compared between the treatments using a linear mixed-effect model with the random effect at individual animal level using R v.3.1 version. The differences between the tumor volumes of mice at the end point were evaluated by Mann-Whitney test.

## Supporting information

Supplemental movie-Cpd1

Supplemental movie-DMSO control

Supplemental movie-Neg4 control

## Acknowledgements

We thank A. Squire and A. Brenzel from the Imaging Center Essen and V. Benes from the Genomics Core Facilities Gene Core at the EMBL Heidelberg; S. Scharfenberg, A. Höwner, A.Cherouny, P. Braß and M. Xiao for their technical support, and Andrew Kossenkov and Patricia Brafford from the Wistar Institute for cDNA microarray analysis and Brian Gabrielli for scientific advice. The results shown are partly based on data from the TCGA Research Network (http://cancergenome.nih.gov/). This work was partly funded by the Hiege Stiftung gegen Hautkrebs, the Monika Kutzner Stiftung, the Deutsche Forschungsgemeinschaft (DFG, German Research Foundation) – RO 3577/3-2, RO 3577/7-1, PA 2376/1-1, BE 1394/12-1, HE 5294/2-1, SCHA 422/17-1, HO 6389/2-1, RE 2857/4-1 (KFO 337), the Australian National Health and Medical Research Council - APP1084893, the National Institutes for Health - PO1 CA114046, P50 CA174523, U54 CA224070, the Excellence Initiative of the German Federal and State governments (Seed Funds programme, project OPSF363), the Dr. Miriam and Sheldon G. Adelson Medical Research Foundation and by grants dedicated to JNRL from the Ministerio de Economia y Competitividad (MINECO; Co-financing with Fondos FEDER) (SAF2016-77241-R), and the Fundación Séneca, the Región de Murcia (FS-RM) (20809/PI/18).

## Author Contributions

HC, AR designed, planned, and evaluated the experiments and wrote the manuscript. HC, BS, SMD, RJJ, SE, AS, FK, OK, KJ, CK, VWR and LK performed the experiments. DP, RV, OV, SJS, QL, XY, FCEV, SSL, JF, SH and IH performed analysis of genome-wide data, statistics and the screening approach. MR, ME, AP, JCB, IH, DR, MK, MH, JNRL, NKH, DS and AR supervised the project. BS, SMD, RJJ, OK, SJS, SG, JNRL and NKH contributed to the design and interpretation of the experiments. All authors were involved in critical revision of the manuscript and approved the final submitted version.

## Competing interests statement

The authors declare no competing financial interests.

## Availability of materials

Except for Tet-On 3G-*KDM5B* and WM3734^Tet3G-shJARID1B^ cell lines, all materials are commercially available. The inducible Tet-On 3G-*KDM5B* and WM3734^Tet3G-shJARID1B^ cell lines are available upon request from the corresponding author [AR] via institutional MTA procedures.

## Notes

### Competing Interest Statement

The authors have declared no competing interest.

