## Supplementary Information for "Persister state-directed transitioning and vulnerability in melanoma"

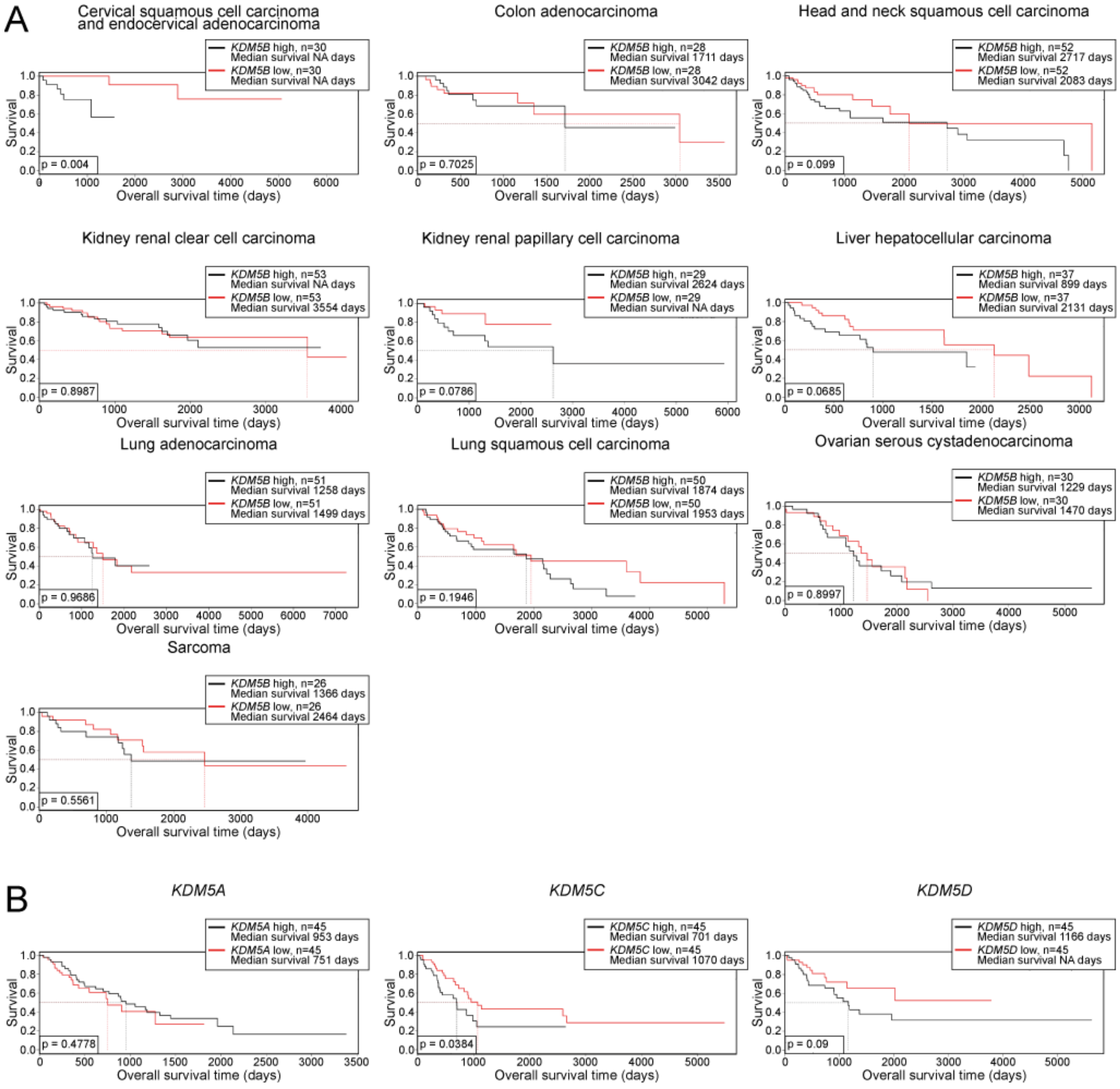


Figure S1

**Figure S1. KDM5 gene-dependent patient overall survival across different cancer entities, Related to Figure 1.** (**A**) Kaplan-Meier survival curves for KDM5B across different cancer entities. Curves were calculated from the TCGA data set based on a 10% gene expression threshold (10% highest expression, black, vs. 10% lowest, red). Graphs were created by the TCGA browser tool v0.9. (**B**) Kaplan-Meier survival curves of melanoma patients for KDM5A, KDM5C, and KDM5D.

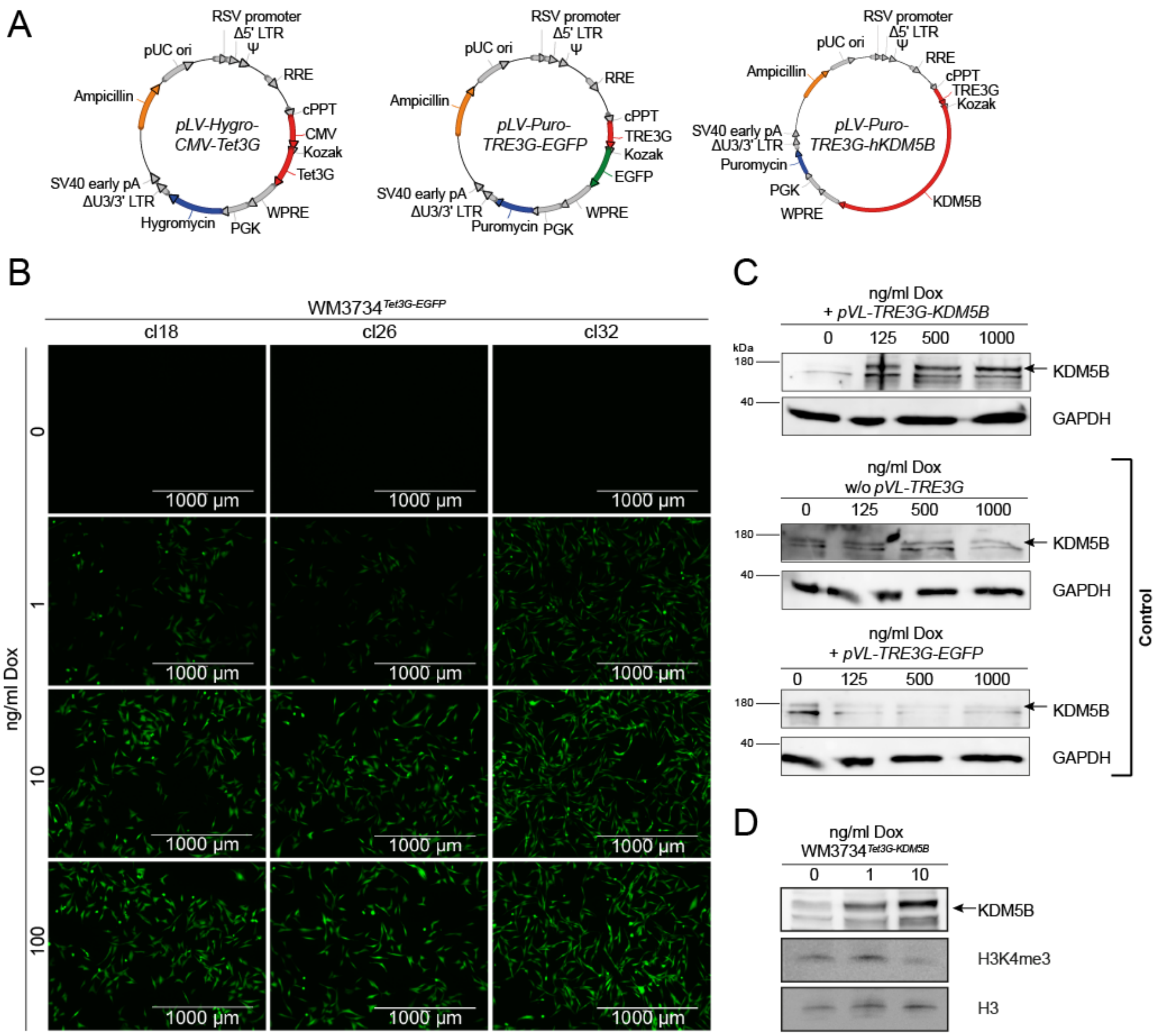


Figure S2

**Figure S2. Establishment of a doxycycline-inducible Tet-On 3G-system for KDM5B protein overexpression, Related to Figure 1.** (**A**) Vector maps of the plasmid encoding the CMV promoter-driven Tet3G (left, pLV-Hygro-CMV-Tet3G), the TRE response vector with an inducible $P_{TRE3G}$ promoter plus the control gene EGFP (middle, pLV-Puro-TRE3G-EGFP), and the TRE response vector with an inducible $P_{TRE3G}$ promoter plus the human KDM5B gene (right, pLV-Puro-TRE3G-hKDM5B, transcript variant 1, NM_001314042.1). (**B**) EGFP protein induction at increasing Dox concentrations in three different clones of stably transduced WM3734$^{Tet3G\text{-}EGFP}$

control cells imaged after 48 h of induction. (**C**) Immunoblotting of KDM5B protein induction in WM3734$^{Tet3G}$ cells transduced with pVL-TRE3G-KDM5B at different Dox levels (top panel). Dox was titrated up to 1000 ng/ml to exclude cell toxicity. Absence of KDM5B protein induction in WM3734$^{Tet3G}$ control cells (middle) and WM3734$^{Tet3G\text{-}EGFP}$ control cells (bottom). (**D**) Representative immunoblots of KDM5B and histone H3K4me3 after 22 days of Dox induction in WM3734$^{Tet3G\text{-}KDM5B}$ cells (n=2).

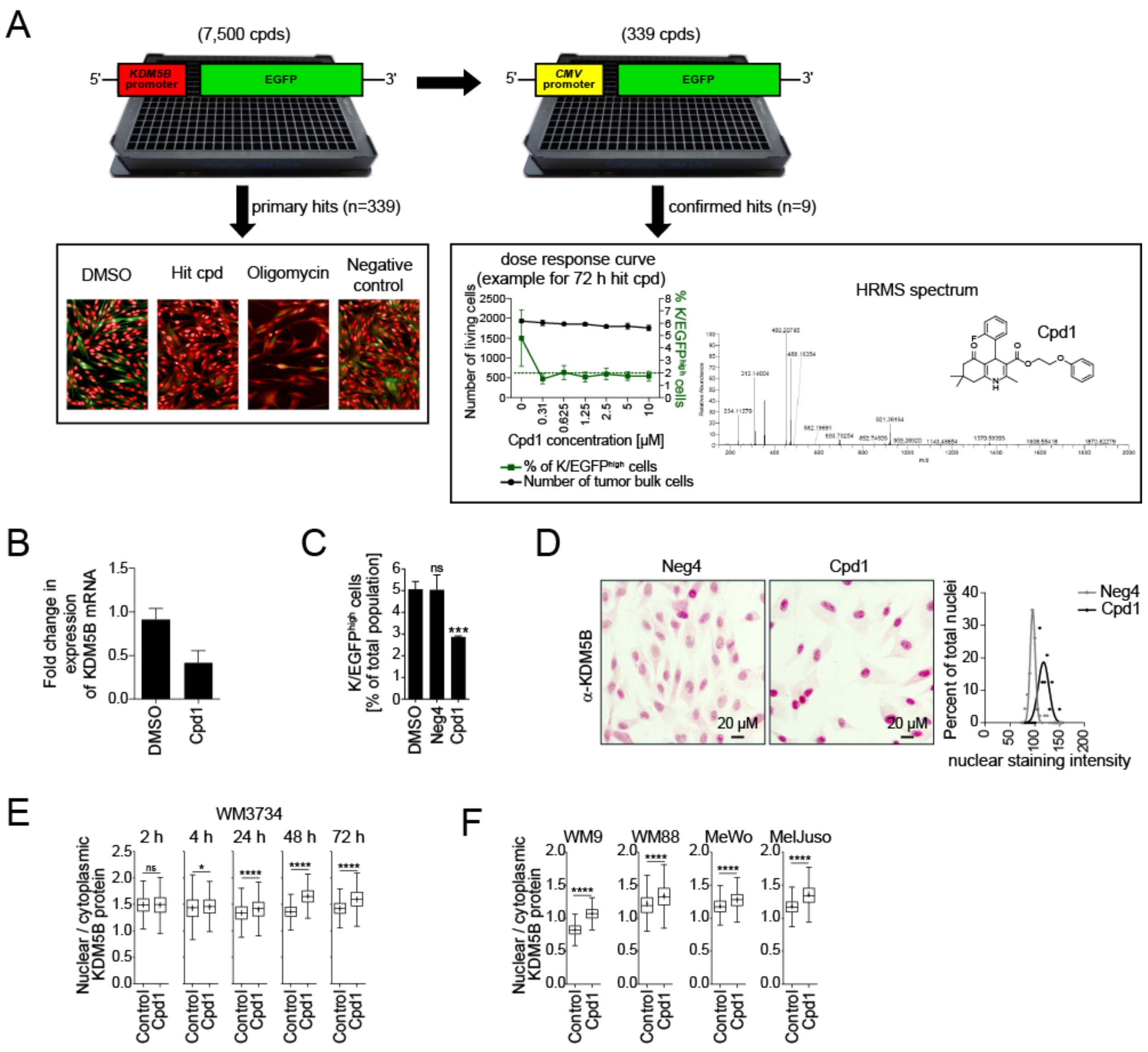


### Figure S3

**Figure S3. Screening for small chemical compounds that modulate KDM5B expression, Related to Figure 1.** (**A**) Schematic of the phenotypic compound screening assay consisting of a primary screen based on the WM3734$^{KDM5Bprom\text{-}EGFP}$ cell model and a counter screen with WM3734$^{CMVprom\text{-}EGPF}$ control cells performed on an Opera High Content Screening system. Dose response curves were done for confirmed hit compounds, as e.g. compound no. 1 (abbreviated

Cpd1). The structural formula and HRMS purity analysis of Cpd1 is shown at the right. For more details, see methods. (**B**) Quantitative PCR detection of endogenous KDM5B mRNA transcripts (all isoforms covered) after 72 h of Cpd1 treatment in WM3734 cells. Shown is one representative experiment out of 4 independent replicates. (**C**) Independent hit validation by flow cytometric detection showed reduced KDM5B-promoter-driven EGFP (K/EGFP) expression in WM3734 cells after 72 h of treatment with the hit compound Cpd1 (10 µM) compared to DMSO and the structural analog compound (Molecular ACCess System (MACCS) similarity of 0.899) Neg4 (10 µM) which lacks comparable activity. Shown are summarized data (mean ±SD) of 3 independent experiments; significance was tested by t-test (***P≤0.001). (**D**) Anti-KDM5B nuclear immunostaining of WM3734 melanoma cells after 72 h of Cpd1 vs. Neg4 treatment (10 µM). Left, representative pictures; right, quantitation shown as normalised frequency distribution of nuclear staining intensity. (**E**) Digital microscopic quantitation of nuclear vs. cytoplasmic KDM5B protein expression in WM3734 cells after Cpd1 treatment or DMSO as control over time and (**F**) across different melanoma cell lines (WM9, WM88, MeWo, MelJuso) after 72 h of treatment. Mean ±SD; *P≤0.05, ****P≤0.0001 by t-test. Box-and-whiskers represent median values and interquartile range; the mean values are plotted as crosses. Shown are results from 2-4 independent replicates with at least 3 quantified images per experiment.

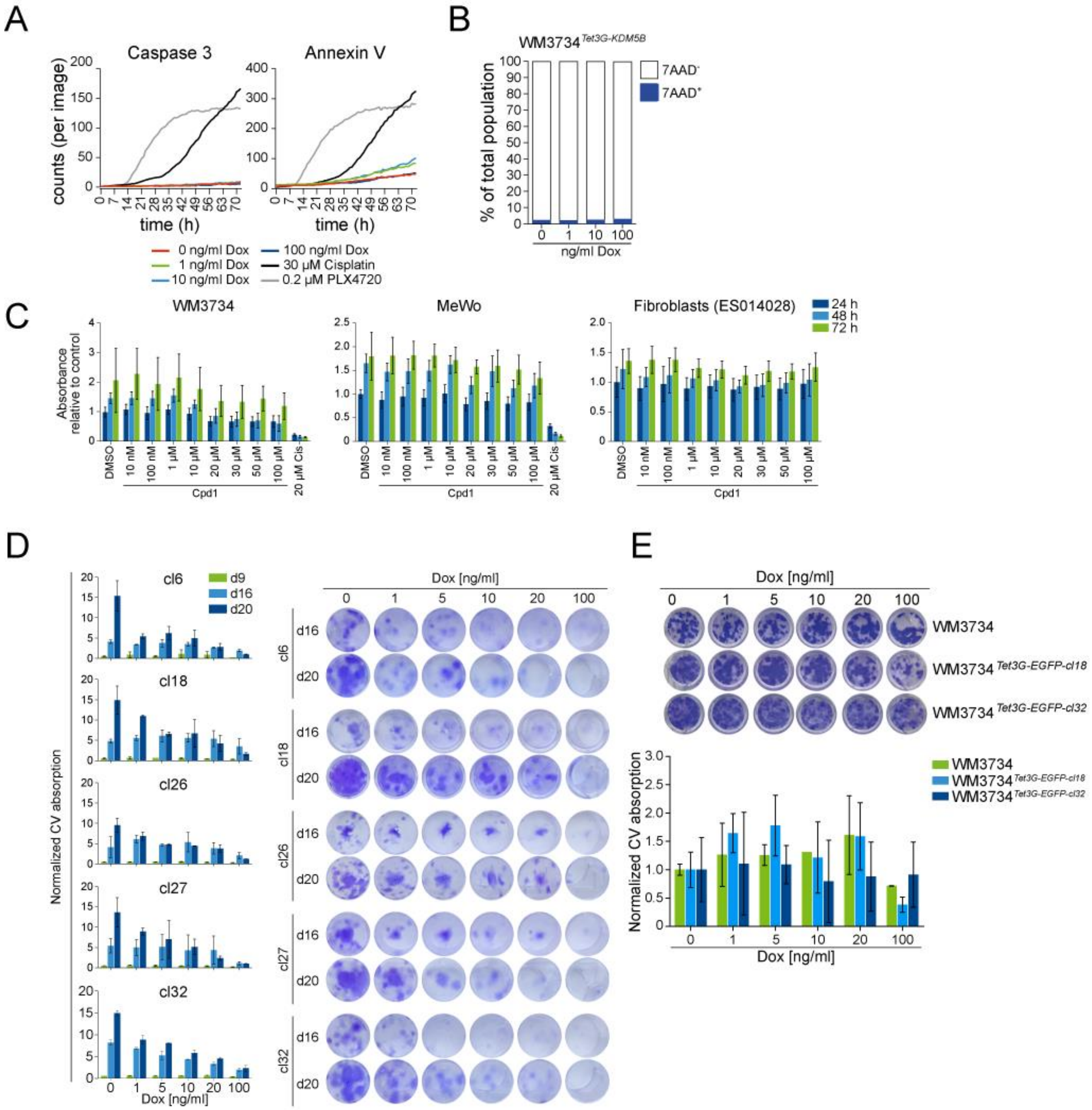
A
Caspase 3
Annexin V
counts (per image)
time (h)
0 ng/ml Dox
1 ng/ml Dox
10 ng/ml Dox
100 ng/ml Dox
30 µM Cisplatin
0.2 µM PLX4720
B
WM3734 Tet3G-KDM5B
% of total population
7AAD-
7AAD+
ng/ml Dox
C
WM3734
MeWo
Fibroblasts (ES014028)
24 h
48 h
72 h
Absorbance relative to control
Cpd1
D
Normalized CV absorption
cl6
cl18
cl26
cl27
cl32
d9
d16
d20
Dox [ng/ml]
E
WM3734
WM3734 Tet3G-EGFP-cl18
WM3734 Tet3G-EGFP-cl32
Normalized CV absorption
Dox [ng/ml]

Figure S4

**Figure S4.** ***In vitro*** **effects of enforced KDM5B expression, Related to Figure 2.** (**A**) Caspase 3 and annexin V detection over 72 h determined by IncuCyte analysis. Cisplatin and PLX4720 were used as positive controls for induction of apoptosis. Shown is one representative experiment out of three independent biological replicates. (**B**) Flow cytometric determination of $7AAD^{+}$ dead cells after 48 h of KDM5B induction by Dox. Shown is one representative experiment out of 4 independent replicates. (**C**) MTT assay after 24, 48, and 72 h of Cpd1 treatment in WM3734 and MeWo melanoma cells compared to primary ES014028 fibroblasts. Data are depicted as mean ±SD. Shown is one representative experiment out of three independent biological replicates. (**D**) Long-term clonogenic growth assay after gradual KDM5B induction over 9, 16, and 20 days with Dox from 5 different $WM3734^{\textit{Tet3G-KDM5B}}$ clones (left, crystal violet quantitation; right, representative pictures from day 16 and 20). Data are shown as mean ±SD. (**E**) Clonogenic long-term growth of $WM3734^{\textit{Tet3G-EGFP}}$ control clones and naïve WM3734 cells after 20 days of Dox exposure (upper panel, representative pictures; lower panel, crystal violet quantitation). Data are shown as mean ±SD.

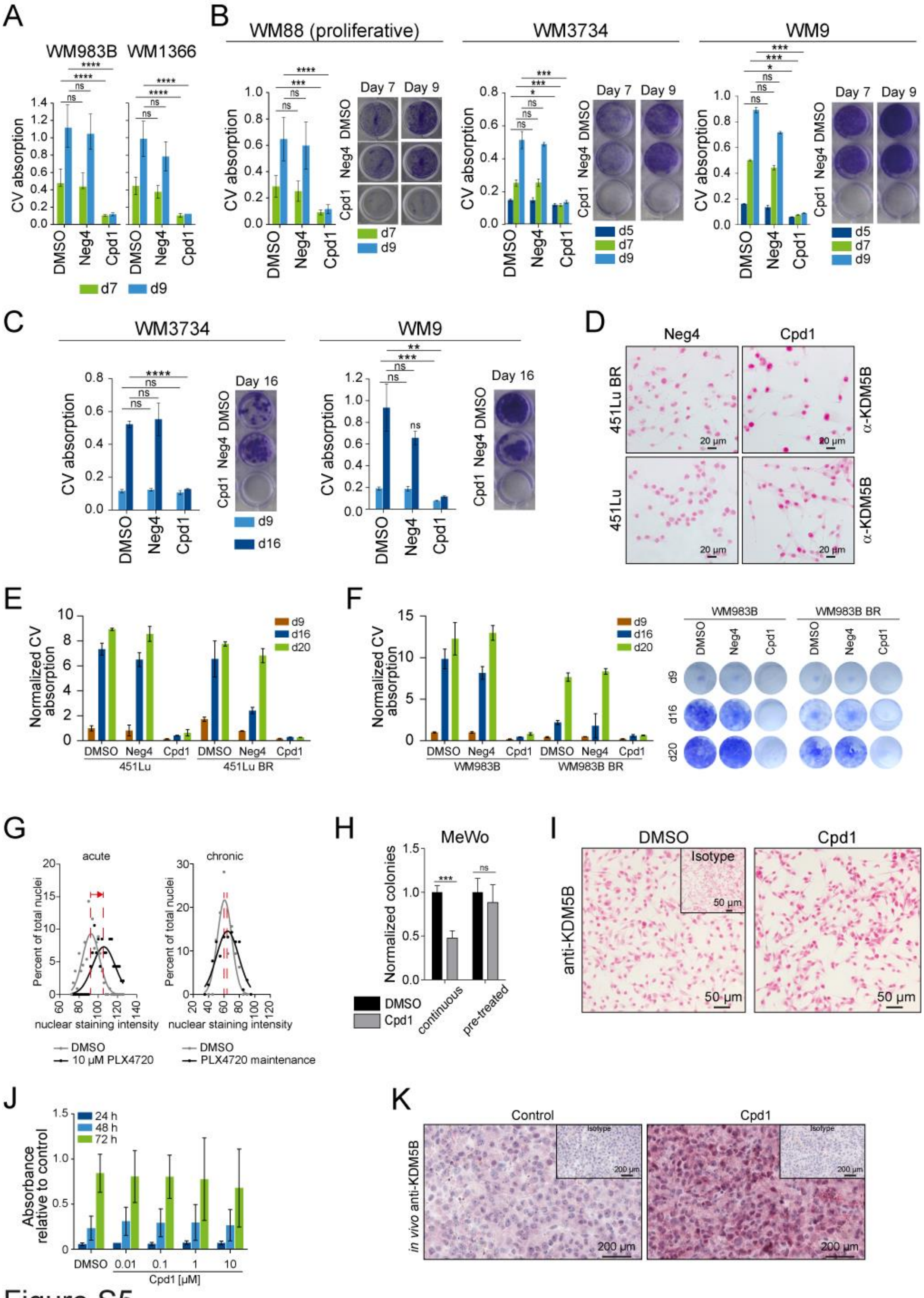
A
WM983B WM1366
CV absorption
DMSO
Neg4
Cpd1
d7
d9
B
WM88 (proliferative)
WM3734
WM9
Day 7
Day 9
d5
C
Day 16
d16
D
Neg4
Cpd1
451Lu BR
451Lu
α-KDM5B
20 µm
E
Normalized CV absorption
d20
F
WM983B
WM983B BR
G
acute
chronic
Percent of total nuclei
nuclear staining intensity
DMSO
10 µM PLX4720
PLX4720 maintenance
H
MeWo
Normalized colonies
continuous
pre-treated
I
Isotype
anti-KDM5B
50 µm
J
Absorbance relative to control
24 h
48 h
72 h
Cpd1 [µM]
K
Control
in vivo anti-KDM5B
200 µm

Figure S5

**Figure S5.** ***In vitro*** **and** ***in vivo*** **effects of enforced KDM5B expression, Related to Figure 2.** (**A**) Quantitation of clonogenic growth assay of Fig. 2E. Shown is one representative experiment out of 3 independent biological replicates (mean ±SD). Significance was assessed by t-test (****P≤0.0001). (**B**) Clonogenic growth assay of WM88, WM3734 and WM9 cells continuously treated over 5, 7 and 9 days with 10 µM of Cpd1 or Neg4 (left, crystal violet quantitation; right representative pictures from day 7 and 9). Shown is one representative experiment out of 3 independent biological replicates (mean ±SD). Significance was assessed by t-test (*P≤0.05, ***P<0.001, ****P≤0.0001). (**C**) Clonogenic growth assay of WM3734 and WM9 cells continuously treated over 9-16 days with 10 µM of Cpd1 or Neg4 (left, crystal violet quantitation (mean ±SD); right representative images from day 16). Significance was determined by t-test (**P≤0.01, ***P<0.001, ****P≤0.0001). (**D**) Representative images of anti-KDM5B immunostaining of chronically PLX4720-resistant 451Lu BR vs. treatment-naïve 451Lu melanoma cells. Cells were treated for 72 h with Cpd1 (10 µM) or Neg4 (10 µM). (**E**) Quantitation of clonogenic growth assay of 451Lu and 451Lu BR cells shown in Fig. 2F. Data are depicted as mean ±SD. (**F**) Clonogenic growth assay of WM983B and PLX4720-resistant WM983B-BR cells treated over 9,16 and 20 days with 10 µM of Cpd1 or Neg4 (left, crystal violet quantitation; right, representative pictures). Data are depicted as mean ±SD. (**G**) Quantitated anti-KDM5B immunostaining shown as normalised frequency distribution of nuclear staining intensity. Treatment-naïve 451Lu melanoma cells were exposed to 10 µM PLX4720 over 72 h and compared to the DMSO control (left panel). Chronically resistant 451LuBR cells were maintained under the presence of 1 µM PLX4720 and compared to naïve 451Lu under DMSO treatment (right panel). (**H**) Soft agar colony formation of MeWo cells under constant Cpd1 treatment or after pre-treatment with Cpd1 for 72 h before seeding. Shown are results of 2 independent experiments with each being performed in triplicate reaction. Data are shown as mean ±SD; ***P≤0.001 by t-test. (**I**) Anti-KDM5B nuclear immunostaining of CM melanoma cells after 72 h of *in vitro* Cpd1 or as control DMSO treatment. (**J**) MTT assay after 24, 48, and 72 h of Cpd1 treatment in CM cells (mean ±SD, n=3). (**K**) Anti-

KDM5B nuclear immunostaining of CM melanoma tumor grafts from Cpd1-treated mice and control mice.

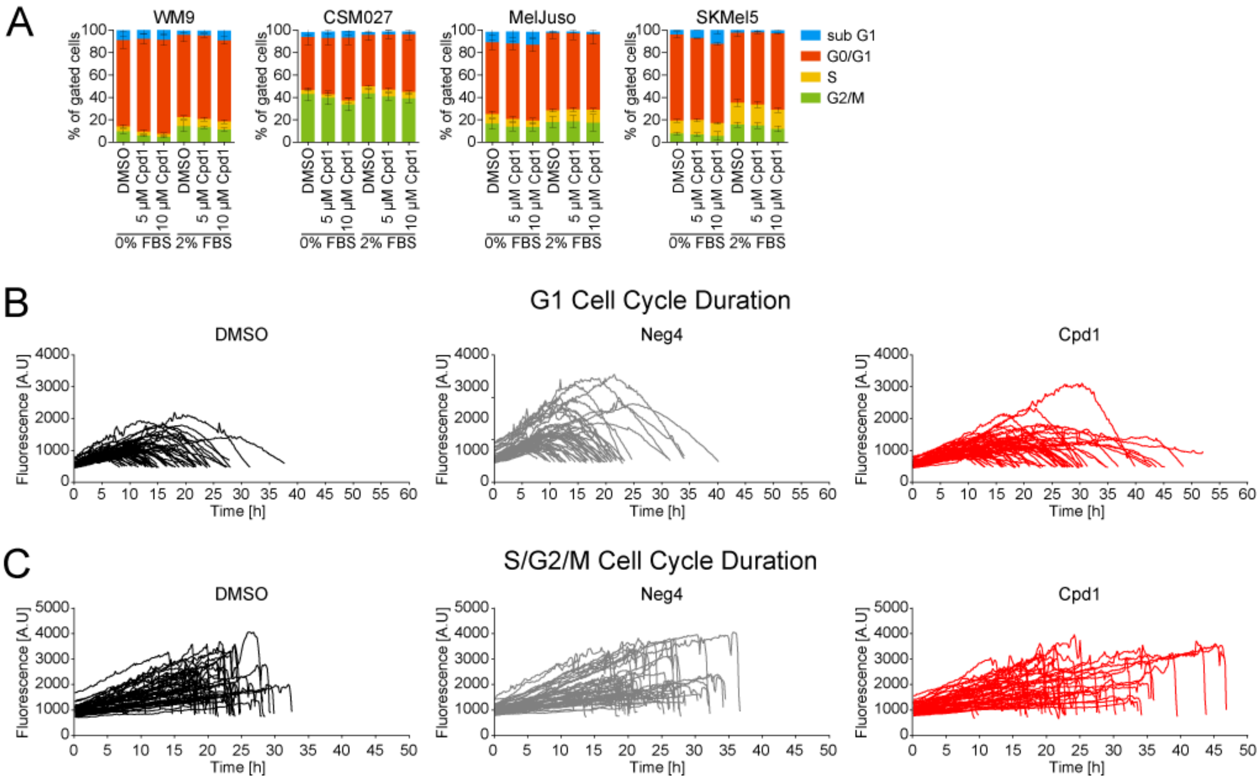


Figure S6

**Figure S6. Effects of enforced KDM5B expression on cell cycle, Related to Figure 3.** (**A**) Propidium iodide flow cytometric cell cycle analysis after 72 h of Cpd1. WM9, CSM027, MelJuso, and SKMel5 cells were either continuously starved (0% FBS) or starved and then released by 2% FBS. Shown are results from 4 independent experiments. Mean ±SD. (**B** and **C**) G1 (**B**) and S/G2/M (**C**) cell cycle duration of single cells (n=49) measured by FUCCI time-lapse imaging after 72 h of treatment with Cpd1 (10 µM) vs. DMSO or Neg4 (10 µM) controls.

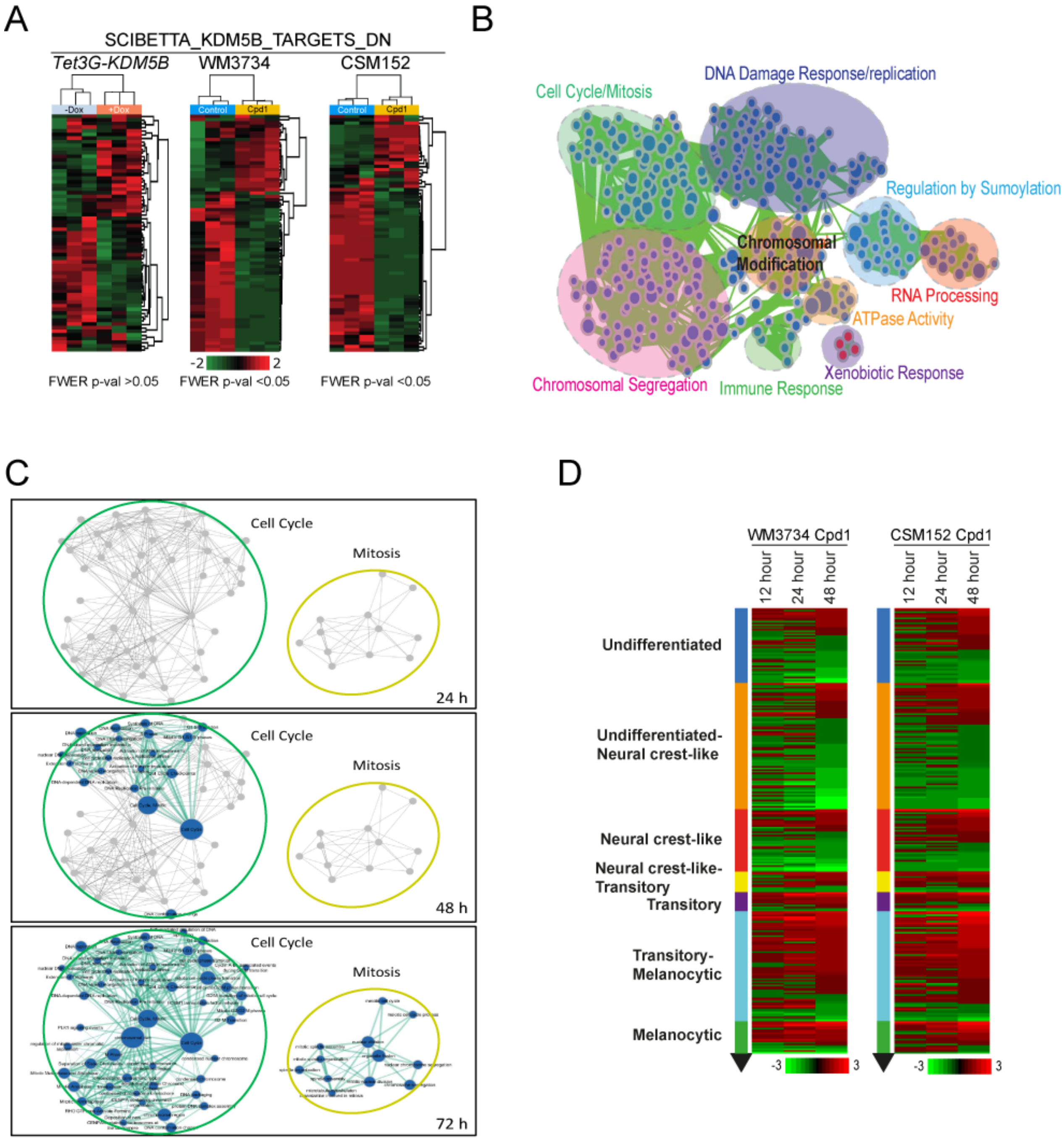


### Figure S7

**Figure S7. Transcriptional shifts as a result of enforced KDM5B expression, Related to Figure 4.** (**A**) Heatmaps of the SCIBETTA_KDM5B_TARGETS_DN (DN='down') motif [1] from WM3734$^{Tet3G\text{-}KDM5B}$ cells after 72 h of Dox treatment compared to corresponding WM3734 cells and short-term cultured CSM152 cells after 72 h of Cpd1 treatment (red, upregulated; green downregulated genes). Significance is indicated by FWER p-val < 0.05. (**B**) Overview of KDM5B-dependent regulatory pathways. Cytoscape enrichment analysis of significantly regulated gene signatures detected in RNAseq in WM3734 and patient-derived, short-term cultured CSM152 cells treated with Cpd1 over 72 h. Red node, enrichment in treatment group; blue node, decrease. The size of the nodes represents the number of genes included. (**C**) Enrichment analysis of cell cycle- and mitosis-controlling transcripts detected in WM3734$^{\textit{Tet3G-KDM5B}}$ cells after 24, 48 and 72 h of Dox treatment. GSEA visualizations for each time point were combined in Cytoscape. Blue nodes represent enrichment in the control group; grey nodes were not yet present at the respective time point, the size of the nodes reflects the number of genes included. (**D**) Heatmap of Tsoi differentiation signature [2] from WM3734 and CSM152 cells after 12 h, 24 h and 48 h Cpd1 treatment ranked by expression at 48 h (red, upregulated; green downregulated genes).

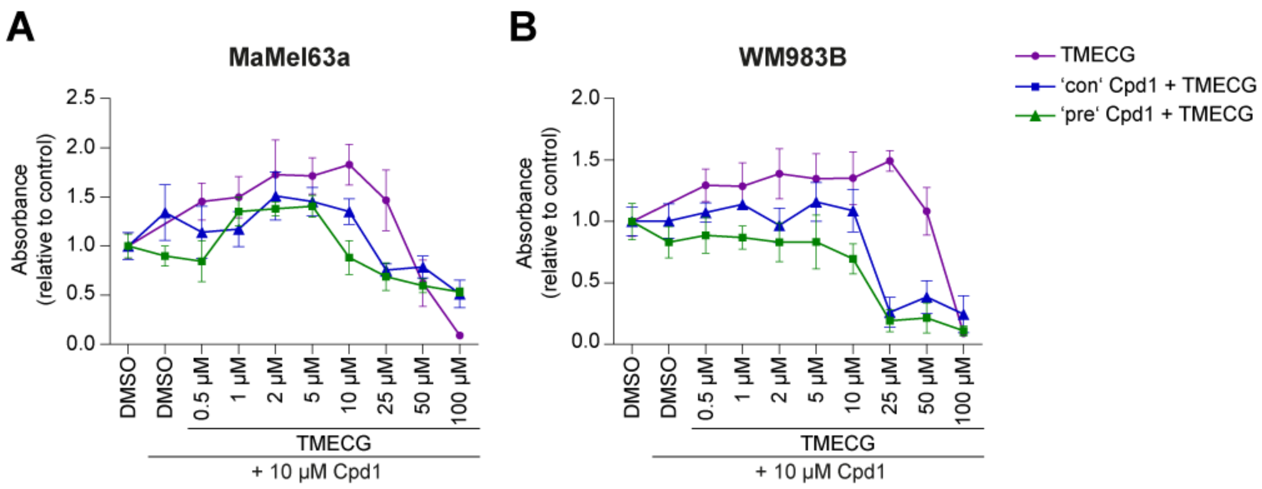


### Figure S8

**Figure S8. Decreased cell viability upon combined Cpd1 plus TMECG treatment, Related to Figure 5.** (**A and B**) MTT cell viability assays of MaMel63a (A) and WM983B (B) cells. TMECG was either concurrently given together with Cpd1 ('con') or added 3 days after Cpd1 pre-treatment ('pre'). Readout was performed after 72 h of TMECG treatment. Data are shown as mean ±SD.

**Table S1. Screening assay results for the selected hit (Cpd1) and control (Neg4) compound. Related to Figure S3.**

| Cmpd Identifier | Structure | Mother plate (WellIndex) | PS % of K/EGFP$^{high}$ cells mean | HC run I % of K/EGFP$^{high}$ cells mean ±SD (n=3) | HC run II % of K/EGFP$^{high}$ mean ±SD (n=2) | HC run III % of K/EGFP$^{high}$ cells mean ±SD (n=3) | Mean |
|---|---|---|---|---|---|---|---|
| 27849_ChemDiv _2191-2790 **(Cpd1)** | 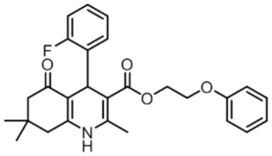 | CBN_000012_D01 (Well I07) | **2.00** | 0.95 ± 0.37 | 1.18 ± 0.24 | 2.27 ± 0.15 | **1.47** |
| 26798_ChemDiv _2732-4408 **(Neg4)** | 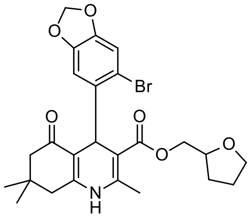 | CBN_000009_D01 (Well N7) | **3.85** | no hit (n.a.) | no hit (n.a.) | no hit (n.a.) | |

PS = Primary Screen
HC = Hit Confirmation

**Table S2. Significantly regulated genes detected by mass spectrometry of WM3734 cells treated with 10 µM Cpd1 for 72 h**. Related to Figure 3.

Please see separately uploaded Table.

**Table S3. Significantly regulated genes detected by RNAseq of cells treated with Cpd1 (WM3734 and CSM152 cells, 10 µM Cpd 1 for 72 h,) or doxycycline (WM3734$^{Tet3G\text{-}KDM5B}$ cells, 10 ng/ml for 24 h, 48 h or 72 h**). Related to Figure 3, 4 and S7.

Please see separately uploaded Table.

**Table S4. Normalized enrichment scores (NES) and false discovery rate (FDR) for the heatmaps of the 'Tsoi et. al differentiation signature' [2] shown in Figure 4C and S7D**. Related to Figure 4 and S7.

| NES | 10 dox vs 0 dox (24 h) | | | WM3734 Cpd1 vs DMSO | | | CSM152 Cpd1 vs DMSO | | |
|---|---|---|---|---|---|---|---|---|---|
| Signature | 24 h | 48 h | 72 h | 12 h | 24 h | 48 h | 12 h | 24 h | 48 h |
| Undifferentiated | 2.13 | -1.52 | -1.50 | -1.38 | -1.35 | -1.93 | -1.58 | -1.14 | -1.76 |
| Undifferentiated-neural crest-like | 1.27 | -2.83 | -2.69 | -1.93 | -1.86 | -2.08 | -2.15 | -1.51 | -1.74 |
| Neural crest-like | -1.73 | -2.67 | -2.60 | -2.26 | -1.93 | -1.79 | -1.17 | -1.64 | -1.27 |
| Neural crest-like-transitory | -1.59 | 1.24 | 1.30 | -1.18 | 1.30 | -1.42 | 1.15 | 1.31 | 1.17 |
| Transitory | 0.87 | 2.17 | 2.16 | -1.04 | 1.02 | -0.89 | 1.13 | 1.89 | 1.66 |
| Transitory-melanocytic | -1.42 | 2.66 | 2.90 | 1.47 | 1.66 | 1.61 | 1.52 | 2.07 | 1.67 |
| Melanocytic | 0.80 | 1.96 | 2.30 | -1.53 | 1.02 | 1.07 | 1.31 | 1.55 | 1.79 |

| FDR | 10 dox vs 0 dox(24 h) | | | WM3734 Cpd1 vs DMSO | | | CSM152 Cpd1 vs DMSO | | |
|---|---|---|---|---|---|---|---|---|---|
| Signature | 24 h | 48 h | 72 h | 12 h | 24 h | 48 h | 12 h | 24 h | 48 h |
| Undifferentiated | 0.00 | 0.03 | 0.03 | 0.11 | 0.09 | 0.00 | 0.03 | 0.28 | 0.03 |
| Undifferentiated-neural crest-like | 0.25 | 0.00 | 0.00 | 0.01 | 0.00 | 0.00 | 0.00 | 0.04 | 0.02 |
| Neural crest-like | 0.02 | 0.00 | 0.00 | 0.00 | 0.00 | 0.01 | 0.24 | 0.03 | 0.14 |
| Neural crest-like-transitory | 0.04 | 0.16 | 0.12 | 0.25 | 0.27 | 0.07 | 0.32 | 0.11 | 0.24 |
| Transitory | 0.93 | 0.00 | 0.00 | 0.39 | 0.55 | 0.64 | 0.26 | 0.00 | 0.01 |
| Transitory-melanocytic | 0.06 | 0.00 | 0.00 | 0.05 | 0.06 | 0.02 | 0.10 | 0.00 | 0.01 |
| Melanocytic | 0.79 | 0.00 | 0.00 | 0.06 | 0.42 | 0.35 | 0.22 | 0.04 | 0.00 |

**Table S5. Significantly regulated genes detected by RNAseq of cells treated with Cpd1 (WM3734 and CSM152 cells, 10 μM for 12 h, 24 h and 48 h)**. Related to Figure S7.

Please see separately uploaded Table.

**Table S6. Primer Sequences**. Related to Figure 1, 3, 5, and S3.

| Name | Species | Sequence (5'-3') | Source |
|---|---|---|---|
| AURKB fwd | human | CATCACACAACGAGACCTATCGCC | |
| AURKB rev | human | GGGTTATGCCTGAGCAGTTTGGAG | |
| KDM5B all isoforms fwd | human | AACAACATGCCAGTGATGGA | [3] |
| KDM5B all isoforms rev | human | TACCAGGTTTTTGGCTCACC | [3] |
| KDM5B transcript variant 1 fwd | human | AACCTCCGCCTCCTAGATTC | |
| KDM5B transcript variant 1 rev | human | CGTTGTCTCCTCGGGTTCTA | |
| KIF4A fwd | human | GAAGAAAACCAAGGCTGAAGGGG | |
| KIF4A rev | human | TGGAATCTCTGTAGGGCACAAAGC | |
| MITF fwd | human | CCGTCTCTCACTGGATTGGT | [4] |
| MITF rev | human | TACTTGGTGGGGTTTTCGAG | [4] |
| SHCBP1 fwd | human | TGTTTGACCAGACAGCCCTTGC | |
| SHCBP1 rev | human | TCATCCTCCTCTTCTTCATCCCAAC | |
| UBE2C fwd | human | GCATCAGAACCAGCTCAACA | |
| UBE2C rev | human | GGTTCTGGCATTTGGAGAAA | |
| 18S fwd | human | CGCCGCTAGAGGTGAAATTC | |
| 18S rev | human | TCTTGGCAAATGCTTTCGC | |
| Myco-SE | human | GGGAGCAAACAGGATTAGATACCC | [5] |
| Myco-AS | human | TGCACCATGTGTCACTCTGTTAACCTC | [5] |

| | | | |
|---|---|---|---|
| DCT fwd | human | AACTGCGAGCGGAAGAAACC,<br>ID 193788638c2 | [6] |
| DCT rev | human | CGTAGTCGGGGTGTACTCTCT,<br>ID 193788638c2 | [6] |
| MITF fwd | human | CCGTCTCTCACTGGATTGGT,<br>ID 8917552a1 | [4] |
| MITF rev | human | TACTTGGTGGGGTTTTCGAG,<br>ID 8917552a1 | [4] |
| MLANA fwd | human | GCTCATCGGCTGTTGGTATT,<br>ID 110625784c2 | [7] |
| MLANA rev | human | TTCTTGTGGGCATCTTCTTG,<br>ID 110625784c2 | [7] |
| TYR fwd | human | GCAAAGCATACCATCAGCTCA | [6] |
| TYR rev | human | GCAGTGCATCCATTGACACAT | [6] |

**Supplemental movies. Live cell imaging of WM3734 cells treated with DMSO, Neg4 (10 µM) or Cpd1 (10 µM) over 5 days. Related to Figure 4.**

Please see separately uploaded supplementary material.
